# The RNA virome of early metazoans sheds light on long-term virus-host relationships

**DOI:** 10.64898/2026.08.30.748148

**Authors:** Ayda Susana Ortiz-Baez, Jonathon C.O. Mifsud, Jasper Schwarz, Sabrina Sadiq, Edward C. Holmes

## Abstract

Ctenophores and placozoans arose early in metazoan evolution and are characterized by traits associated with key aspects of animal evolution. Despite the evolutionary significance of ctenophores and placozoans, their RNA viromes are poorly understood. To determine the diversity and evolution of RNA virome in these organisms, particularly whether the viruses present with these ancient host lineages similarly occupy basal phylogenetic positions, we analysed publicly available transcriptome data from the Sequence Read Archive (SRA). Accordingly, we identified 26 putative novel viruses classified into 11 virus groups, including members of the families *Flaviviridae* and *Chuviridae*. The novel viruses clustered with those previously identified in vertebrates, invertebrates, plants and fungi. Notably, some virus sequences within the *Flaviviridae*, *Chuviridae*, *Lispiviridae* and *Marnaviridae* were highly divergent, branching deeply relative to their closest known relatives or forming distinct lineages, in some cases suggesting a divergence early in metazoan evolution. In contrast, viruses within the *Birnaviridae*, *Endornaviridae*, *Mymonaviridae*, *Narnaviridae*, *Phasmaviridae*, *Orthomyxoviridae*, *Orthototiviridae*, and some viruses within the *Picornavirales*, exhibited patterns consistent with more recent diversification and host jumping. In addition, RNA viruses were detected across multiple species and tissues within the Ctenophora (including whole organisms and embryos) and Placozoa, expanding their host range and highlighting a largely uncharacterized diversity. Together, these findings expand the known diversity and host range of several virus groups, and shed light on virus evolution in early metazoans, demonstrating both host jumping within aquatic environments and virus host-associations that may span the entirety of animal evolution.

## Introduction

Ctenophores (Ctenophora, also known as comb jellies) and placozoans (Placozoa) are two of the earliest diverging lineages within the Metazoa (i.e., animals) and hence of considerable evolutionary importance. These organisms possess anatomical traits that mark key transitions in the evolution of animal body plans, including tissue organization (e.g., neural tissue) [1–3], body symmetry [4, 5], digestive structures [6], and other aspects of biological organization [7, 8]. The early origin and biological uniqueness of ctenophores and placozoans also raises important questions about the viruses that may be associated with these hosts, particularly whether they similarly occupy basal positions in viral phylogenies and the nature of virus-host interactions over long evolutionary timescales [9].

Although the Ctenophora and Placozoa are evidently ancient animal groups, the identity of the earliest-diverging metazoan lineage remains contentious, particularly whether sponges (Porifera) [10–14] or ctenophores represent the first branch from the last common ancestor of animals, likely dating back to some point in the Neoproterozoic era ∼1000–541 million years ago (MYA) [15, 16]. Discovered in the 18^th^ century [17], ctenophores exist in polar to tropical marine environments [4] distributed throughout the water column and even occurring at depths that reach the hadal zone (>6,000 m), and are represented by over 200 fossil and living species [4, 18]. These marine predators feed on such prey as zooplankton, copepods, cladocerans, larvae, and might also engage in cannibalism [19, 20]. Notably, they are characterized by biradial symmetry and structures such as a through-gut, eight comb rows equipped with locomotory cilia, tentacles, mesoglea and the aboral organ which functions as a sensory and gravity structure [4]. Ctenophores also exhibit muscle fibres and a non-centralized nervous system composed of a subepithelial nerve net, which concentrate in the aboral organ and polar fields [21, 22].

Placozoans are small marine animals with an undifferentiated bauplan, typically 1-10 mm in diameter, nerveless and lacking any apparent symmetry [23, 24]. These organisms originated during the Tonian period, approximately 800–750 MYA [2], and while only a few species have been recently recognized, numerous remain unclassified or referred as “haplotypes” [25, 26]. Placozoan bodies are composed of approximately 20,000 cells, comprising nine cell types with specialized roles, including gland, lipophile, fibres and epithelial peptidergic cells, with the latter involved in locomotion and paracrine signalling (i.e., neuron-like cells) [2]. Notably, placozoans possess some of the smallest genomes among metazoans (at ∼100 Mb), while their mitochondrial genomes are among the largest, reaching up to ∼43 Kb [23, 25]. Reproduction in this group includes asexual and sexual strategies, in which vegetative budding and fission are the most common mechanisms under laboratory conditions [23, 26]. Ecologically, placozoans are the smallest free-living animals, commonly found in shallow and calm waters up to ∼20 m depth [27]. Feeding on organic detritus, bacteria, algae, protozoans, and biofilms via transepithelial cytophagy [24], they are primarily found in tropics and subtropics, with some records in more temperate waters [23, 28, 29].

Although our understanding of the biology of ctenophores and placozoans continues to expand [30], little is known about the viruses that infect these animals [9, 31–34]. Large-scale analyses of viral diversity across metazoan phyla tentatively suggest that ctenophores and placozoans may harbour less diversity than other invertebrate phyla [32]. However, most studies have not targeted these early-diverging lineages directly through metagenomic sequencing [33, 34]. Although previous studies have identified flavivirus and orthomyxovirus sequences in ctenophores and other animal phyla [33, 34], suggesting that these viral families have an ancient evolutionary origin, it is unclear whether viruses present in the Ctenophora and Placozoa represent deeply divergent lineages, thereby reflecting long-term co-divergence throughout metazoan evolution, or result from repeated host-jumping events within a marine environment.

Herein, we mined publicly available transcriptome sequencing data from ctenophores and placozoans to reveal more of their RNA virome diversity and to determine whether the viruses present in these early metazoans form basal branches within established viral families or have been acquired more recently through cross-species transmission. We also provide insights into the broader evolution of tissue tropism and host-virus associations in metazoans.

## Methods

### SRA screening and data collection

Publicly available sequencing data from the Ctenophora (taxid 10197) and Placozoa (taxid 10226) were retrieved from the NCBI Sequence Read Archive (SRA) between October-December 2025. The search included filters to retain only transcriptomic data (“RNA-Seq” [Assay type] AND “Illumina/DNBSeq” [Platform] AND “Paired/Single” [Library Layout]). Sequencing reads were downloaded using Kingfisher v.0.4.1 [35]. Accordingly, 211 sequencing libraries were assembled comprising 203 paired-end libraries from ctenophores and 28 from placozoans.

### Raw data processing, contig assembly and abundance estimation

Raw reads were initially assessed for quality using FastQC v0.12.1 [36]. Quality trimming was performed with Trimmomatic v0.35 [37] using an average Phred quality threshold of 20, followed by a second reassessment with FastQC. For the Third Party Annotation (TPA) assembly, contigs were *de novo* assembled using MEGAHIT v1.29 with default parameter settings [38]. The relative abundance of virus contigs within metatranscriptomes was calculated as the number of transcripts per million (TPM) using RSEM v1.3.0 [39]. Viruses present at a frequency of <0.1% of the highest abundance observed for a given virus contig across libraries were assumed to represent index-hopping and excluded from all subsequent analyses.

### Contig annotation and taxonomic profiling

The assembled contigs were initially screened for sequences of likely viral origin by comparisons against the (i) RNA dependant RNA polymerase database (RdRp) [40] and (ii) the protein Reference Viral Database (RVDB-prot clustered) v31 [41]. Subsequently, putative viral contigs were annotated through searches against the non-redundant (nr) NCBI database (https://blast.ncbi.nlm.nih.gov/Blast.cgi) with the E-value threshold set to ≤1E-4 using DIAMOND v.2.1.10 [42]. These contigs were then compared against the nucleotide (nt) NCBI database using an E-value of ≤ 1E-10. Similarly, viral contigs were subjected to open reading frame (ORF) prediction using the standard genetic code in ORFfinder (https://www.ncbi.nlm.nih.gov/orffinder/) and annotated for domains and motifs using the Conserved Domain Database (CCD) [43] and InterProScan [44] with the full suite of integrated databases. For placozoan sequences, ORF prediction was also performed using the Mold, Protozoan and Coelenterate Mitochondrial code (which identified no additional viruses). Accordingly, sequences were classified as putative viruses if they were ≥ 600 nt in length, produced a hit against the databases described above, and contained conserved viral genes or domains, such as the RdRp and capsid proteins. Finally, the overall sequence composition of libraries (i.e., to confirm that they were associated with the Ctenophora and Placozoa) was assessed using Kraken2 v.2.17.1 (core-nt database v.11-2025) [45] and reports were visualized using Pavian v.1.0 [46]. Taxonomic assignment was performed using TaxonKit [47].

### Protein structural prediction

Glycoprotein sequences were obtained from the closest known relatives to those viruses identified here in the BLASTp searches as well as for reference sequences in the respective virus groups. In the case of the flavi-like viruses, the boundaries of the envelope protein ectodomain (E) were determined using the structure and sequence of tick-borne encephalitis virus (PDB: 7QRF) alongside predicted E structures from the orthoflaviviruses and flavi-like viruses taken from Mifsud et al. 2024 [48]. Protein structures were then predicted using the AlphaFold3 web server (https://alphafoldserver.com/) (release 2025.07.05) [49], with default settings apart from disabling template usage. The five predictions for each model were examined through Predicted Aligned Error (PAE) and predicted local distance difference test scores, and in all cases the top ranked prediction by AlphaFold3 based on overall structural confidence scores was selected and used for all downstream analyses. All structural visualisations were prepared using UCSF ChimeraX v.1.10 [50]. To establish whether the predicted structures shared structural similarity to existing virus membrane fusogens, the predicted structures were queried against extracted monomers of experimental PDB structures representing the three membrane fusion protein classes (I,II,III) using Foldseek v.10-941cd33 [51] easy-search with non-default parameters (--num-iterations 3 and -- exhaustive-search) (Supplementary Table 1). Foldseek e-value scores against each representative structure were visualised using ggplot2 package v.4.0.1 [52] in R v.4.5.0 [53].

### Phylogenetic analyses and protein structure prediction

RdRp amino acid sequences from putative viruses were aligned with those from their closest known relatives in the BLAST searches, and additional reference sequences in the respective virus groups, using the L-INS-i algorithm in MAFFT v.7.525 [54]. Phylogenetic relationships among these sequences were inferred using the maximum likelihood (ML) method available in IQ-TREE v.2.3.6 [55] with ModelFinder [56] (-m MFP) employed to find the best-fit model of amino acid substitution. Node support was estimated with 1000 ultrafast bootstrap [57](UFBoot) replicates and the Shimodaira-Hasegawa approximate likelihood ratio test [58] (SH-aLRT), with topological confidence set as SH-aLRT >= 80% and UFboot >= 95%, respectively.

To incorporate protein structural information into our phylogenetic inference we implemented a variation of the 3DiPhy approach described previously [59, 60]. Accordingly, we converted our predicted full protein structures to 3Di sequences with the Foldseek (release 10) ‘structureto3didescriptor’ option and used MAFFT with Foldseek 3Di character substitution matrix and parameters “--genafpair --maxiterate 1000” to infer structural, 3Di sequence alignments of the E1, helicase and RdRp protein sets. We used trimAl v.1.5.1 [61] with the automated “-gappyout” option to create trimmed versions of the 3Di multiple sequence alignments (MSAs). In addition to the 3Di character alignments, we replaced the 3Di characters with the protein amino acid residues in both the complete and trimmed versions of the MSAs. This resulted in four MSAs (3Di, trimmed 3Di, amino acid, trimmed amino acid) for each protein. Modelfinder implemented in IQ-TREE 3 v.3.0.1 [62] was again used to determine the best substitution model for each alignment. Various models were tested, including a custom 3Di substitution calculated from Foldseek’s 3Di substitution cost matrix model [60] (i.e., -mset Blosum62, Dayhoff, DCMut, JTT, JTTDCMut, LG, Poisson, Poisson+FQ, Poisson, PMB, WAG, EX2, EX3, EHO, EX_EHO, 3DI), with amino acid frequency options specified by the protein matrix, or empirical base frequencies (-mfreq FU,F) and among-site variations models (-mrate E,G,R) equal, gamma, or FreeRate. The selected substitution models for all alignments are shown in Supplementary Table 2. Phylogenetic trees based on each MSA were again inferred using the ML approach in IQ-TREE3 under each corresponding best-fit substitution model, with node support assessed using 1000 ultra-fast bootstraps (UFBoot) and SH-aLRT replicates. Phylogenetic inference was performed using both the 3Di character and amino acid sequences by combining the corresponding pairs of 3Di and amino acid MSAs and performing edge-linked partition model phylogenetic analysis in IQ-TREE 3, where each partition was allowed its own evolutionary rate and best-fit substitution model (flag -p) [63].

### Workflow and code availability

Computational analyses were performed on the Gadi (National Computing Infrastructure, Australia) and Setonix (Pawsey Supercomputing Centre) high-performance computing systems. The analytical procedure and code are implemented in the Batch Artemis SRA Miner pipeline v. v1.0.4 (https://github.com/JonathonMifsud/BatchArtemisSRAMiner/tree/v.1.0.4), with adaptations made as needed.

## Results

### Overview of the early metazoan data set and associated RNA viruses

To investigate the RNA viromes of early-diverging metazoan lineages, we mined transcriptomic data corresponding to 203 bulk RNA sequencing libraries from ctenophores and 28 from placozoans, including 12 single-cell sequencing libraries (Figure 1A-B). Overall, the sequencing libraries comprised multiple species of ctenophores and placozoans, with ctenophore libraries spanning a range of different tissues and structures (Figure 1C-E). Collectively, the analyses comprised 745 GB of sequencing data. A total of 6,515,788 contigs were assembled, of which ∼0.02% originated from RNA viruses (Figure 2A). Viral contigs were identified across several ctenophore and placozoan species (Figure 2B, from a variety of tissue types, partial, and whole animal samples (Figure 2C).

**Figure 1.**
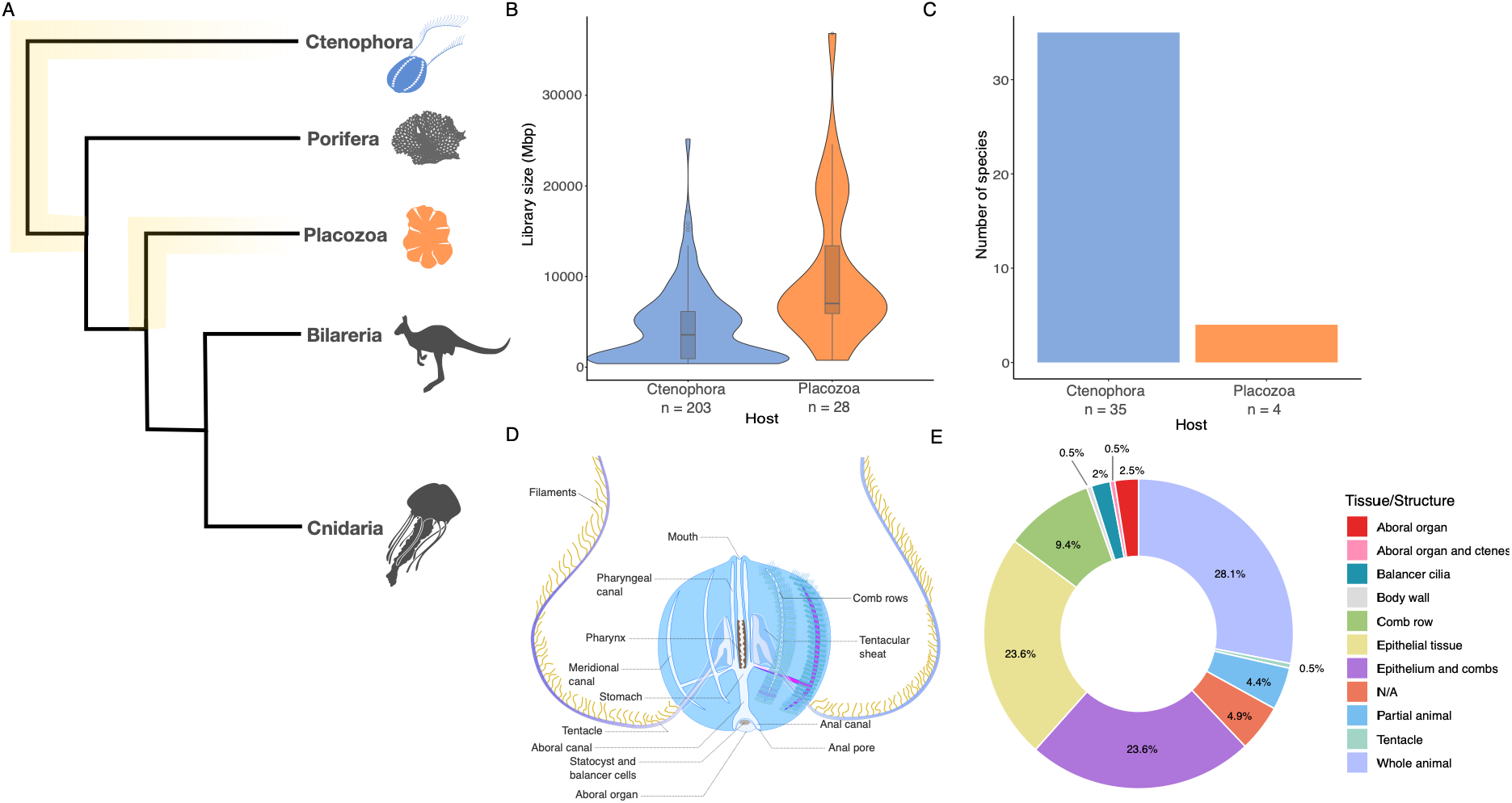
Overview of the SRA library data set used for virus discovery in ctenophores and placozoans. (A) Schematic phylogenetic tree of the Metazoa, with ctenophores placed as the sister lineage to all other animals (although this is debated). (B) Distribution of sequencing library sizes (total reads) per host (C) Number of species represented in the sequencing libraries of each host. (D) Representation of ctenophore body structures from which sequencing libraries were generated. Figure adapted from Wikipedia under CC BY-SA 4.0 https://creativecommons.org/licenses/by-sa/4.0/. (E) Proportional representation of tissues/structures across sequencing libraries of ctenophores.

**Figure 2.**
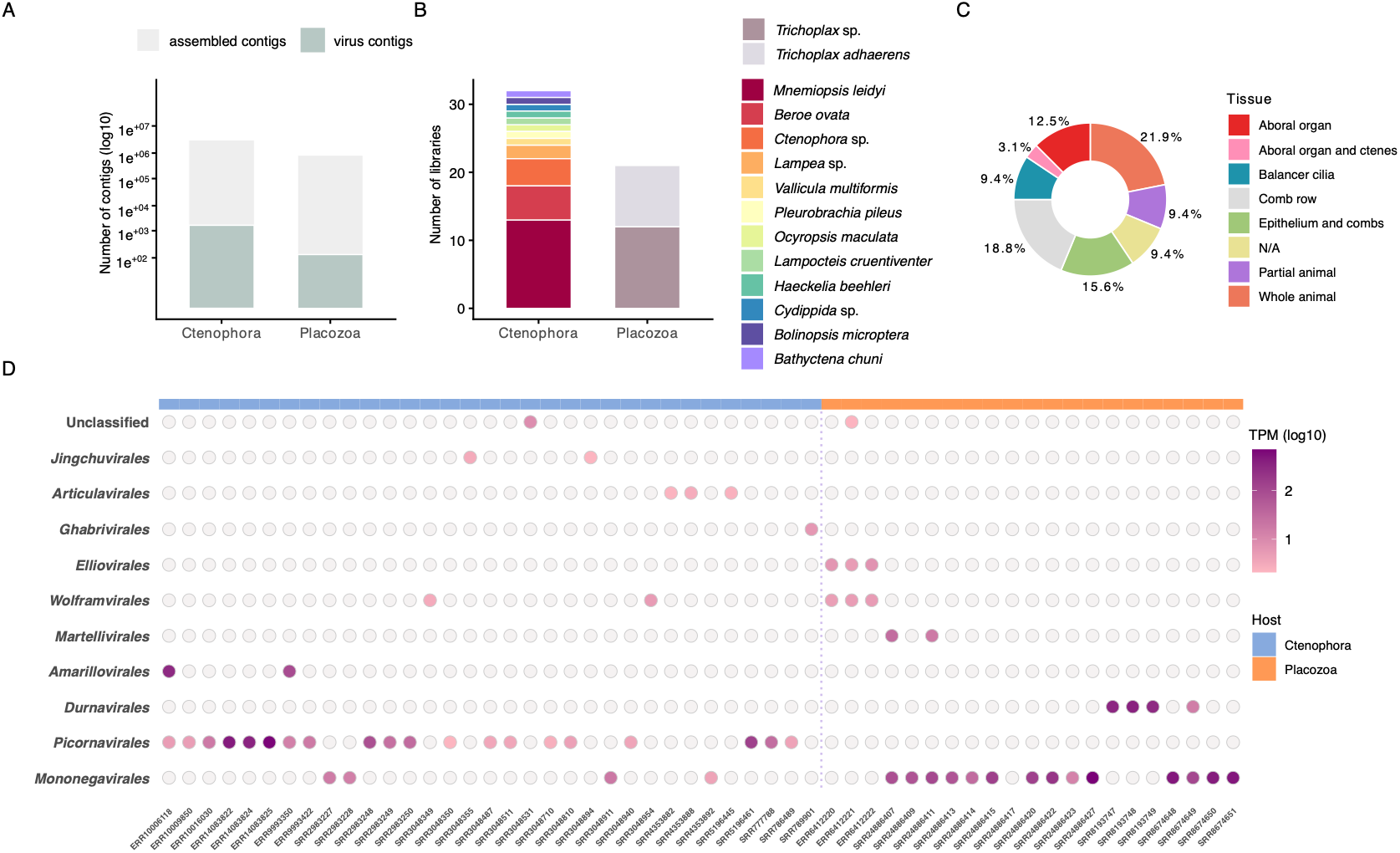
Summary of SRA sequencing libraries containing viral sequences. (A) Proportion of assembled contigs classified as viral for each host. (B) Host species composition of libraries containing viral reads. (C) Distribution of tissue types in ctenophore libraries with viral reads. (D) Overview of viral abundance by virus group across SRA sequencing libraries, quantified as the number of transcripts per million (TPM). Libraries are grouped by host in the top bar (Ctenophora in orange on the left, Placozoa in blue on the right).

From these data we identified 26 viruses belonging to 11 virus groups: order *Amarillovirales* (family *Flaviviridae*), *Jingchuvirales* (*Chuviridae*), *Picornavirales* (*Dicistroviridae* and unclassified), *Mononegavirales* (*Lispiviridae*, *Mymonaviridae and Rhabdoviridae*), *Durnavirales* (*Fusariviridae*), *Martellivirales* (*Endornaviridae*), *Elliovirales* (*Phasmaviridae*), *Articulavirales* (*Orthomyxoviridae*), *Ghabrivirales* (*Orthototiviridae*), *Wolframvirales* (*Narnaviridae*) and unclassified (*Birnaviridae*) (Table 1, Figures 2D, Figures 3-5). Notably, host composition analyses showed that libraries were dominated by unclassified sequences (70%–93%), suggesting either low-quality sequences or an underrepresentation of placozoans and ctenophores in the reference database, which may potentially impact taxonomic classification. Among the classified sequences, host-derived sequences from Ctenophora and Placozoa accounted for most cases (Placozoa: 41.83% – 99.41%; Ctenophora: 13.71% – 99.51%), indicating likely limited cross-contamination with non-host sources.

**Figure 3.**
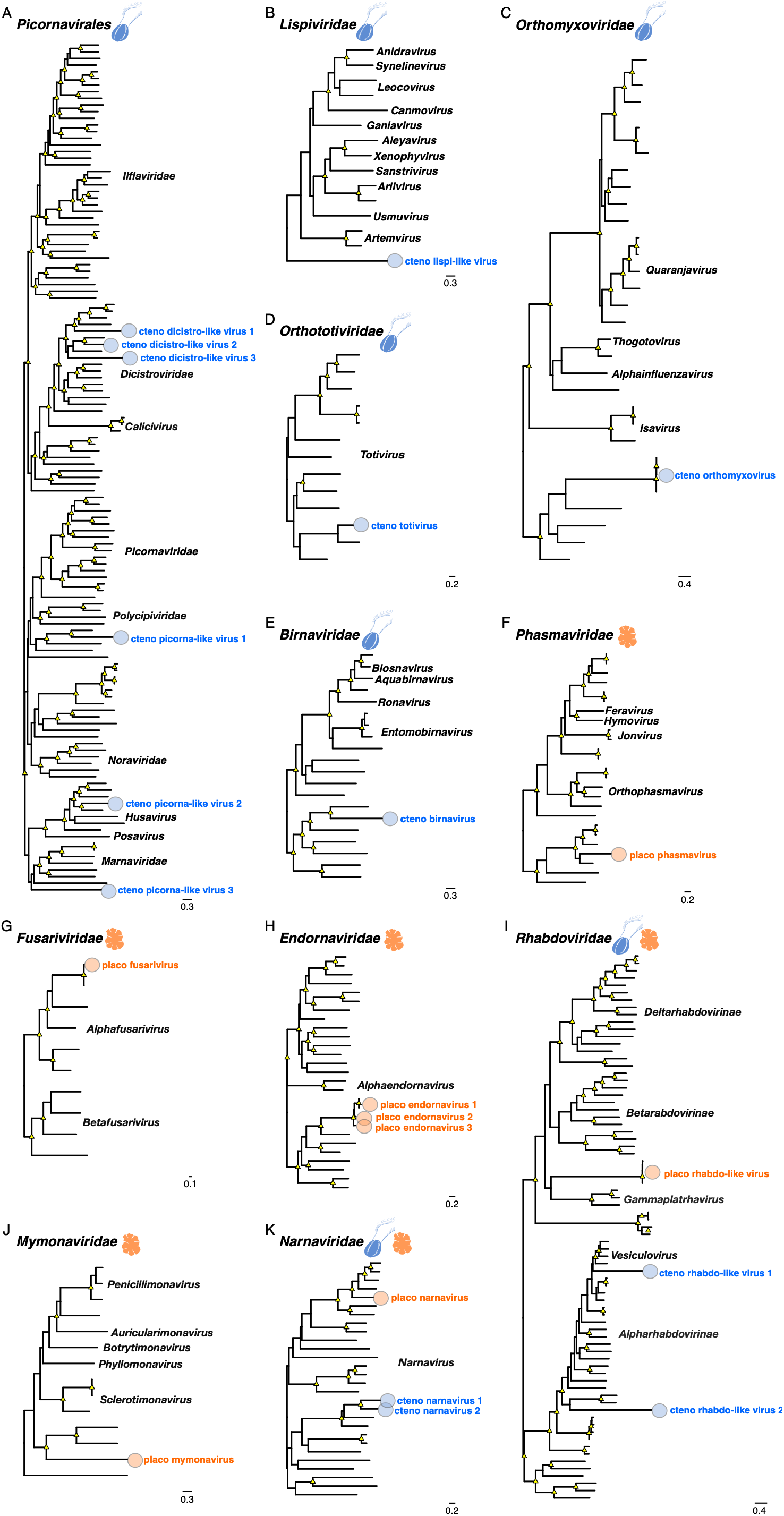
ML phylogenetic trees for viral groups identified in ctenophores and placozoans. (A) *Picornavirales*, (B) *Lispiviridae*, (C) *Orthomyxoviridae*, (D) *Orthototiviridae*, (E) *Birnaviridae*, (F) *Phasmaviridae*, (G) *Fusaviridae*, (H) *Endornaviridae*, (I) *Rhabdoviridae*, (J) *Mymonaviridae* and (K) *Narnaviridae*. All trees were estimated using the amino acid sequence of the RdRp and midpoint rooted for purpose of clarity. Animal silhouettes indicate the host library source. The viruses identified in this study are highlighted in blue or orange for ctenophore and placozoan hosts, respectively. The scale bar indicates the number of substitutions per site. Well-supported nodes are denoted with yellow triangle shapes (SH-aLRT ≥ 80 % and UFboot ≥ 95 %). Full phylogenies with all tips labelled are shown in Supplementary Figure 1.

**Table 1.** RNA viruses identified in ctenophores and placozoans, with matches in the NR/NCBI database based on RdRp sequences.

| Host taxon | Provisional Virus Name | SRA library | Length | Best hit in NCBI/nr | % Identity | E-value | Classification |
| --- | --- | --- | --- | --- | --- | --- | --- |
| **Ctenophora** |  |  |  |  |  |  |  |
| *Ocyropsis maculata* | Cteno chuvirus 1 | SRR3048355 | 899 | QHX39772.1 RNA-dependent RNA polymerase [Herr Frank virus 1] | 50 | 3.94E-12 | *Chuviridae*;<br>*Piscichuvirus* |
| *Haekelia beehleri* | Cteno chuvirus 2 | SRR3048894 | 2336 | WLN26260.1 MAG: L protein, partial [African cichlid piscichuvirus] | 40.2 | 5.42E-174 | *Chuviridae*;<br>*Piscichuvirus* |
| *Beroe ovata* | Cteno orthoflavivirus | ERR10006118 | 11652 | QJU12405.1 polyprotein [Salmon flavivirus] | 32.6 | 4.11E-242 | *Flaviviridae* |
| *Beroe ovata* | Cteno orthoflavivirus | ERR9993350 | 11413 | QJU12405.1 polyprotein [Salmon flavivirus] | 32.6 | 2.82E-242 | *Flaviviridae* |
| *Mnemiopsis leidyi* | Cteno rhabdo-like virus 1 | SRR4353892 | 667 | YP_009505531.1 polymerase [Carajas virus] | 41.8 | 3.81E-50 | *Rhabdoviridae*;<br>*Vesiculovirus* |
| *Mnemiopsis leidyi* | Cteno rhabdo-like virus 2 | SRR4353892 | 720 | AQX45764.2 polymerase protein, partial [Benxi bat virus] | 40.8 | 1.71E-54 | *Rhabdoviridae* |
| *Ctenophora* sp. | Cteno lispi-like virus | SRR3048911 | 7774 | YP_009288955.1 RNA-dependent RNA polymerase [Sanxia water strider virus 4] | 22.9 | 4.34E-25 | *Lispiviridae*;<br>*Sanstrivirus* |
| *Mnemiopsis leidyi* | Cteno orthomyxovirus | SRR4353882 | 916 | UDL13962.1 MAG: RNA-dependent RNA polymerase subunit PB1 [Xiangshan orthomyxo-like virus] | 24.5 | 1.46E-05 | *Orthomyxoviridae* |
| *Mnemiopsis leidyi* | Cteno orthomyxovirus | SRR4353888 | 1061 | UMO75720.1 MAG: polymerase PB1 [Xinjiang sediment orthomyxo-like virus 1] | 24.2 | 2.13E-13 | *Orthomyxoviridae* |
| *Mnemiopsis leidyi* | Cteno dicistro-like virus 1 | SRR5196461 | 9010 | YP_009336613.1 hypothetical protein 1 [Wenling picorna-like virus 3] | 33.6 | 2.83E-105 | *Picornavirales* |
| *Lampea* sp. | Cteno dicistro-like virus 2 | SRR9162937 | 10189 | YP_009336690.1 hypothetical protein 1 [Wenling crustacean virus 2] | 39.3 | 0 | *Picornavirales* |
| *Ctenophora* sp. | Cteno dicistro-like virus 3 | SRR3048940 | 6088 | QUS52706.1 polyprotein [Mute swan feces associated picorna-like virus 10] | 26.7 | 1.05E-103 | *Picornavirales* |
| *Cydippida* sp. | Cteno picorna-like virus 1 | SRR777788 | 3557 | WWZ85695.1 RNA-dependent RNA polymerase [Penaeus vannamei picorna-like virus 4] | 27.4 | 2.34E-26 | *Picornavirales* |
| *Lampea* sp. | Cteno picorna-like virus 2 | SRR3048810 | 4237 | XHA87699.1 MAG: putative viral coat protein [Skokie dicistro-like virus] | 36.4 | 1.72E-229 | *Picornavirales* |
| *Beroe ovata* | Cteno picorna-like virus 3 | ERR10016030 | 6033 | WPR17502.1 MAG: non-structural polyprotein, partial [Mite picorna-like virus 3] | 38.8 | 4.98E-11 | *Picornavirales* |
| **Ctenophora; microptera** | Cteno birnavirus | SRR3048531 | 3636 | WPR17539.1 MAG: VP1 protein, partial [Spider birna-like virus] | 26 | 1.85E-15 | *Birnaviridae* |
| *Pleurobrachia pileus* | Cteno totivirus | SRR789901 | 3061 | WZH59755.1 MAG: RNA-dependent RNA polymerase [Ripugrir virus] | 42.1 | 3.02E-176 | *Orthototiviridae* |
| *Lampea lactea* | Cteno narnavirus 1 | SRR3048349 | 1209 | ASM94100.1 putative RNA-dependent RNA polymerase, partial [Barns Ness serrated wrack narna-like virus 4] | 33.4 | 6.80e-47 | *Narnaviridae* |
| *Ctenophora* sp. | Cteno narnavirus 2 | SRR3048954 | 2723 | WZH58335.1 MAG: RNA-dependent RNA polymerase, partial [Chrocakinb virus] | 33.8 | 1.62e-62 | *Narnaviridae* |
| **Placozoa** |  |  |  |  |  |  |  |
| *Trichoplax adhaerens* | Placo rhabdo-like virus | SRR24886417 | 13061 | UHK03110.1 MAG: RNA-dependent RNA polymerase [Sanya conocephalus maculatus rhabdovirus 1] | 22.6 | 7.45E-74 | *Rhabdoviridae* |
| *Trichoplax adhaerens* | Placo rhabdo-like virus | SRR24886415 | 9120 | UUG74104.1 MAG: RNA-dependent RNA polymerase [XiangYun mono-chu-like virus 4] | 32.7 | 1.93E-51 | *Rhabdoviridae* |
| *Trichoplax* sp. H2 | Placo rhabdo-like virus | SRR24886422 | 3257 | UDL13993.1 MAG: RNA dependent RNA polymerase [Xiangshan rhabdo-like virus 3] | 27.5 | 9.05E-68 | *Rhabdoviridae* |
| *Trichoplax* sp. H2 | Placo rhabdo-like virus | SRR24886420 | 964 | WWB07779.1 RNA-dependent RNA polymerase [Ixeris denticulata-associated rhabdovirus] | 36.8 | 4.26E-42 | *Rhabdoviridae* |
| *Trichoplax* sp. H2 | Placo phasmavirus | ERR6412220 | 5659 | WIL00279.1 MAG: putative RNA-dependent RNA polymerase [Ditton virus] | 33.5 | 6.42E-301 | *Phasmaviridae* |
| *Placozoa* sp. H4 | Placo fusarivirus | SRR8193749 | 5988 | YP_010799569.1 RNA-dependent RNA polymerase [Zymoseptoria tritici fusarivirus 1] | 100 | 0 | *Fusariviridae* |
| *Trichoplax adhaerens* | Placo mymonavirus 1 | SRR24886413 | 7131 | QQO58800.1 RNA-dependent RNA polymerase [Plasmopara viticola associated mononega-like virus 1] | 22.6 | 4.99E-15 | *Mymonaviridae* |
| *Trichoplax* sp. H2 | Placo endornavirus 1 | SRR24886417 | 12026 | QYF50052.1 MAG: polyprotein, partial [Sichuan alphaendornavirus 2] | 41.9 | 1.17E-94 | *Endornaviridae; Alphaendornavirus* |
| *Trichoplax* sp. H2 | Placo endornavirus 1 | SRR24886407 | 987 | BCL84886.1 polyprotein [Phytophthora endornavirus 2] | 44.1 | 5.58E-70 | *Endornaviridae* |
| *Trichoplax* sp. H2 | Placo endornavirus 2 | SRR24886411 | 1222 | BAK52155.1 polyprotein [Bell pepper alphaendornavirus] | 42.9 | 2.92E-89 | *Endornaviridae; Alphaendornavirus* |
| ***Trichoplax* sp. H2** | Placo endornavirus 3 | SRR24886407 | 1286 | YP_009305414.1 polyprotein [Winged bean alphaendornavirus 1] | 37.1 | 1.86E-20 | *Endornaviridae;*<br>*Alphaendornavirus* |
| ***Trichoplax* sp. H2** | Placo narnavirus | ERR6412221 | 3185 | WQM87268.1 putative RdRP [Vo narna-like virus] | 39.7 | 2.13E-232 | *Narnaviridae* |

### RNA virus diversity and evolution in ctenophores and placozoans

To investigate whether the evolutionary history of RNA viruses in early metazoans mirrors aspects of animal phylogeny by occupying divergent topological positions, we inferred phylogenetic trees for each virus group. Interestingly, only narnaviruses and rhabdoviruses were identified in both ctenophores and placozoans, with the newly discovered viruses distributed across multiple clades within each virus family (Figure 2, Figure 3). Specifically, the newly discovered narnaviruses most often grouped with viral sequences from environmental metagenomic samples associated with fungi (or protozoa), exhibiting up to 40% identity to the closest hits in the NCBI non-redundant (nr) database. Similarly, the novel rhabdoviruses were divergent from known virus sequences (23 – 42%) (Table 1), and clustered with sequences found in diverse hosts, including plants and insects. Notably, many of the viruses identified within the *Rhabdoviridae* displayed long branches and low support values, indicating high divergence and limited resolution in their inferred relationships (Figure 3).

Unsurprisingly, as they contributed most libraries for host screening, the ctenophore libraries contained most virus groups identified in this study. These virus sequences were associated with a range of tissues (including aboral organ, comb rows, embryos) across at least 11 ctenophore species (Figure 2B-D). Screening of ctenophore transcriptomes revealed 17 novel RNA viruses. Viral sequences representing the *Picornavirales*, *Orthomyxoviridae*, *Lispiviridae*, *Birnaviridae* and *Orthototiviridae* were identified in libraries from at least seven ctenophore species (Table 1, Figure 3A-E, Supplementary Figure 1). In several instances these viruses exhibited limited amino acid sequence identity (26 – 42%) with viruses associated with invertebrate hosts and environmental samples (e.g., riverbank sediment). As a consequence, taxonomic assignment and host association could not be confidently assigned (Table 1). As a case in point, the novel cteno totivirus found in *Pleurobrachia pileus* was related to a totivirus identified in riverbank sediment (42.1% amino acid identity) (Figure 3D). While the known host range of totiviruses is currently limited to fungi and protozoans, their potential detection in ctenophores points to a broader host spectrum including animals. Similarly, many of the novel viral sequences represented previously unrecognized lineages that occupied early-diverging positions within their respective virus groups (e.g., *Picornavirales* and *Lispiviridae*). For instance, the novel cteno lispi-like virus was the most basal branch within the *Lispiviridae* (Figure 3B), compatible with the early evolutionary divergence of the ctenophores under a model of virus-host co-divergence. Lispiviruses have been detected in a range of hosts including arthropods, nematodes, mammals and birds. That this virus was found in the comb row of an unclassified ctenophore in the bathypelagic zone raises questions about the mechanisms underlying persistence and viral transmission within this specialized epithelial structure and under such extreme conditions. Among the novel picornaviruses, cteno picorna-like virus 3 was identified in the comb row of *Beroe ovata* and occupied a basal position with respect to the *Marnaviridae*, compatible virus-host co-divergence, although this placement was weakly supported (Figure 3A, Supplementary Figure 1). The remaining picornaviruses did not fall in basal phylogenetic positions, instead forming distinct lineages and grouping with invertebrate-associated viruses (i.e., the *Dicistroviridae* and unclassified), suggesting host-switching events involving invertebrate hosts, although an environmental or dietary host origin cannot be entirely excluded. Similarly, the novel ctenophore birnavirus clustered with viruses derived from invertebrate and environmental metagenomes (Figure 3E), indicating phylogenetic incongruence between viral and host phylogenies likely as a result of a history of host jumping. Also of note was that we identified *Mnemiopsis leidyi* orthomyxo-like virus in two SRA libraries corresponding to whole-animal embryos collected following fertilization. In phylogenetic analysis, this virus represented a divergent sister lineage to an orthomyxovirus found in *Hydra oligactis* (Cnidaria, Hydrozoa), and grouped with other viruses found in invertebrates and environmental-derived samples in a pattern that is more consistent with cross-species transmission (and supporting a previous analysis of orthomyxoviruses in ctenophores (Figure 3C) [33].

We also identified 12 putative RNA viruses present in single-cell sequencing libraries from Placozoa (i.e., *Trichoplax adhaerens* and *Trichoplax* spp.), which argues against cross-contamination from other eukaryotic hosts. These newly discovered viruses from Placozoa were classified within the families *Phasmaviridae, Endornaviridae*, *Narnaviridae*, *Rhabdoviridae* and *Mymonaviridae* (Table 1) and formed distinct phylogenetic lineages within their respective clades (Figure 3F-K, Supplementary Figure 1). For instance, the novel placozoan endornaviruses formed a distinct clade, indicating a substantial and uncharacterized diversity within this group (Figure 3H), and expands the host range of this group from plants, fungi and oomycetes. Notably, the novel placo rhabdo-like virus and placo mymonavirus formed deep lineages relative to their sister taxa (Figure 3I-J), yet neither occupied a basal position within their respective families as expected under virus-host co-divergence (although the phylogenetic position of placo rhabdo-like virus was uncertain reflected in low node support). Overall, the novel placo mymonavirus and placo rhabdo-like virus were related to those identified in plant and fungal metagenomes, sharing 23–44% sequence identity with their closest matches in the NCBI database (Table 1). Interestingly, we also recovered a fusavirus sequence that was identical in the RdRp region (1491 aa) to Zymoseptoria tritici fusavirus 1 (Figure 3G), which was detected in the fungal wheat pathogen *Zymoseptoria tritici*. This finding is consistent with the detection of fungi, which comprised approximately 17–18% of the library composition.

### Flaviviruses and Chuviruses in Ctenophores

Of particular note was that the ctenophore transcriptomes also contained three viruses from the *Flaviviridae* and *Chuviridae* that exhibited 33–50% amino acid identity in the RdRp to known vertebrate-associated viruses of the same families present in the NCBI non-redundant (nr) database (Table 1). Notably, a novel flavivirus was identified in the aboral organ of *Beroe ovata* that was most closely related (∼89% amino acid identity in the RdRp over a 115 amino acid region) to cigar comb jelly flavi−like virus, previously identified in the cigar comb jelly *Beroe forskalii* [34], and hence indicative of a ctenophore clade sister to the genus *Orthoflavivirus* (*Flaviviridae*). Interestingly, this clade in turn formed a well-supported monophyletic group with salmon flavivirus and African cichlid flavi-like virus, found in the brain and the lower pharyngeal jaw of fish, respectively, and that fell as a sister-group to the orthoflaviviruses (Figure 4A).

**Figure 4.**
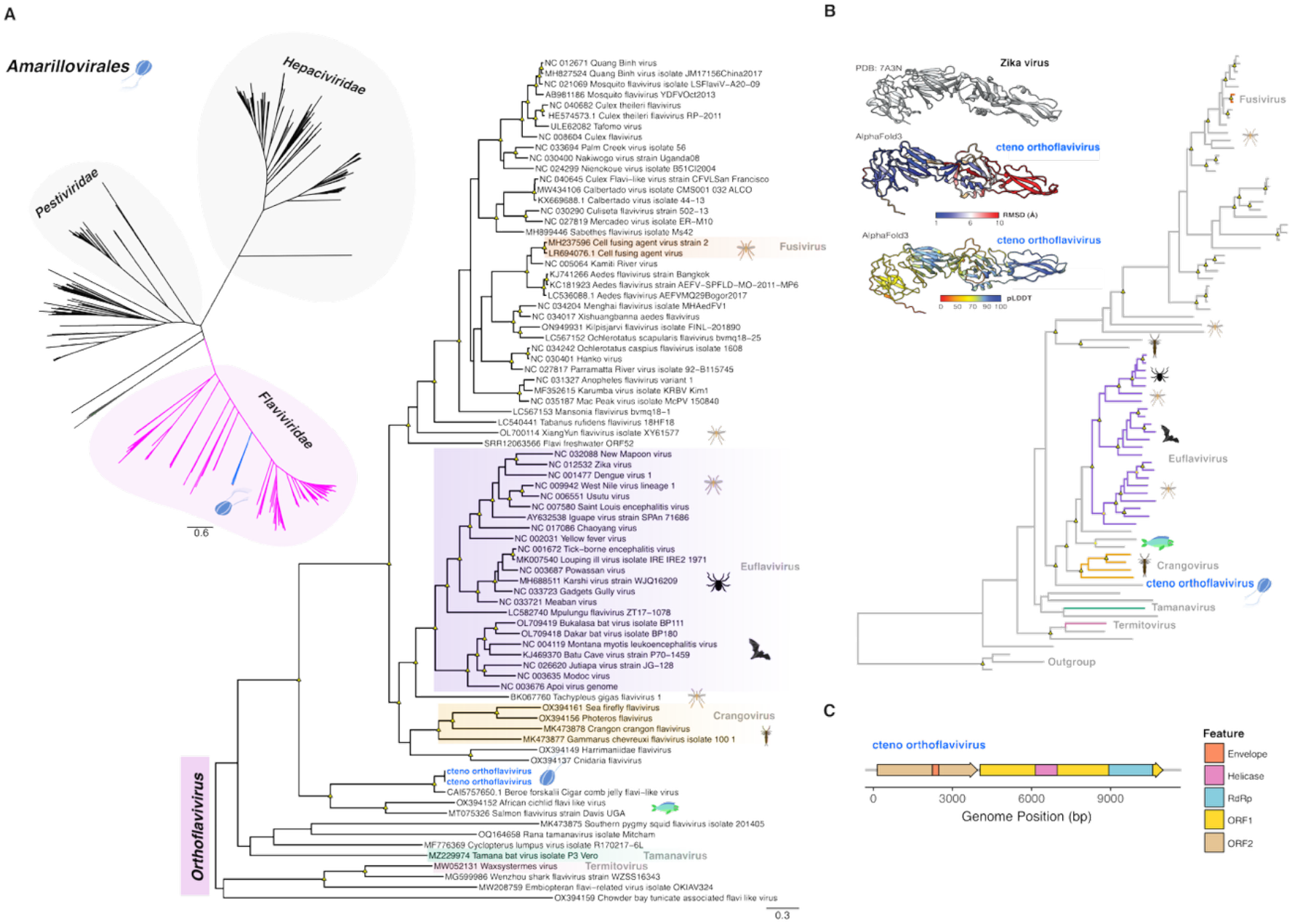
Phylogenetic relationships of the novel orthoflavi-like virus identified in Ctenophora. (A) *Left panel*: unrooted tree showing the relationships among the *Amarillovirales*. *Right panel*: phylogenetic placement of the cteno orthoflavi-like virus within the genus *Orthoflavivirus*. Recognized subgenera within the *Flaviviridae* are labelled where applicable. The virus sequences identified in this study are indicated in bold blue text and represent the same virus. Animal silhouettes denote the most common host for the relevant clades. Newly discovered viruses are indicated by jelly comb silhouettes. The tree is midpoint rooted for purpose of clarity. Both ML phylogenies were estimated based on RdRp amino acid sequences, and the scale bar indicates the number of substitutions per site. (B) *Left panel*: Reference crystal structure of the *Orthoflavivirus* (i.e., Zika virus, PDB 7A3N) glycoprotein and the AlphaFold3-predicted model for the cteno orthoflavi-like virus glycoprotein. The predicted structure is coloured based on the pLDDT and RMSD scores as indicated in the bars. *Right panel*: Phylogenetic tree of the glycoprotein based on a partitioned model combining 3Di and amino acid alignments. Clades are labelled to match the tree in panel A. The scale bar indicates the number of 3Di and amino acid character substitutions per site. The tree is rooted using nairoviruses as the outgroup. Unclassified viruses are shown in grey. In all trees, well-supported nodes are denoted with yellow triangle shapes (SH-aLRT ≥ 80 % and UFboot ≥ 95 %). (C) Schematic depiction of the ORFs and domains recovered for the novel cteno orthoflavi-like virus [48].

Interestingly, the phylogenetic placement of the clade containing salmon flavivirus and African cichlid flavi-like virus varied depending on the methods and the proteins used for phylogenetic analysis [48, 64]. To better characterise the ctenophore flaviviruses and their relationship to the orthoflavi-like viruses across both structural and non-structural proteins, we analysed the E glycoprotein of the ctenophore flavivirus. Amino acid identity between the ctenophore flavivirus E glycoprotein and its closest relative was low (Salmon flavivirus, 27.36% identity). As such, we sought structural information to provide additional evolutionary resolution. Modelling of the ctenophore flavivirus E glycoprotein of the ctenophore flaviviruses using AlphaFold3 revealed a predicted structure broadly consistent with experimentally determined structures of prototypical orthoflavivirus class II fusion proteins (Figure 4B, Supplementary Figure 2, Supplementary Table 1). Specifically, the predicted structure contained a putative fusion loop with the conserved sequence motif characteristic of orthoflavivirus class II fusion proteins (average pLDDT = 93, residues 1,586-1,624), as well as a putative precursor membrane protein (prM, residues 755-1,288) characteristic of orthoflaviviruses and some orthoflavi-like viruses. Together, these features support the phylogenetic placement of this virus within this genus (Figure 4). Due to the high level of sequence divergence across flavivirus glycoproteins, we next estimated structurally informed glycoprotein phylogenies (Figure 4B-C). The first, based on a structurally informed amino acid alignment placed the ctenophore flavivirus in a clade with salmon flavivirus and African cichlid flavi-like virus, was consistent with the RdRp phylogeny. However, the position of this clade differed substantially from the RdRp phylogeny, instead forming a sister group to the euflaviviruses separate from the insect-specific flaviviruses (including the fusiviruses). Additional phylogenies were inferred using 3Di structural characters or a joint partition model of 3Di and amino acid characters (Supplementary Figure 3). Although the separation of the euflaviviruses and fusiviruses was consistent across all of the glycoprotein phylogenies, the position of the ctenophore flavivirus varied. Specifically, in the 3Di phylogeny the ctenophore flavivirus formed a sister group to tamana-termite- and fusi-like viruses, albeit with poor bootstrap support. The joint tree combining 3Di and amino acids was more consistent with the structurally informed amino acid phylogeny, placing the ctenophore flavivirus basal to the euflaviviruses (Figure 4). In this case, the ctenophore flavivirus was basal to the salmon flavivirus and African cichlid flavi-like virus.

The Ctenophora pischu-like viruses (*Chuviridae*) newly identified in this study were present in the whole-body and comb rows of the species *Haeckelia beehleri* and *Ocyropsis maculata*, respectively, clustering together in the phylogenetic analysis yet with only ∼57% amino acid identity among them (Figure 5A), and exhibiting limited similarity with African cichlid piscichuvirus and Herr Frank virus 1 (∼40–50% amino acid identity in the RdRp) in BLAST searches (Table 1). Protein structure prediction revealed that the glycoprotein of novel cteno chuvirus 2 had high probability hits to several class III fusion proteins including Suid alphaherpesvirus 1 (Figure 5B-C, Supplementary Table 1 and Supplementary Figure 4). These Ctenophora pischuvirus sequences fell as a sister lineage to a clade dominated by viruses previously identified in fish, reptile and crustaceans (i.e., decapods), within which likely vertebrate-associated chuviruses have been detected across a range of tissues including the brain, liver, and gills (Figure 5A). Notably, this clade also includes a freshwater macrophyte chuvirus (Freshwater macrophyte associated chu-like virus 1) that moves to a more basal, but less supported, topological position in the partitioned structural phylogenetic analysis of the glycoprotein (Figure 5A-B, Supplementary Figure 5). Although only the 3Di analysis recovered a closer relationship with turtle neural viruses, support across all three phylogenetic analyses remained low (Supplementary Figure 5).

**Figure 5.**
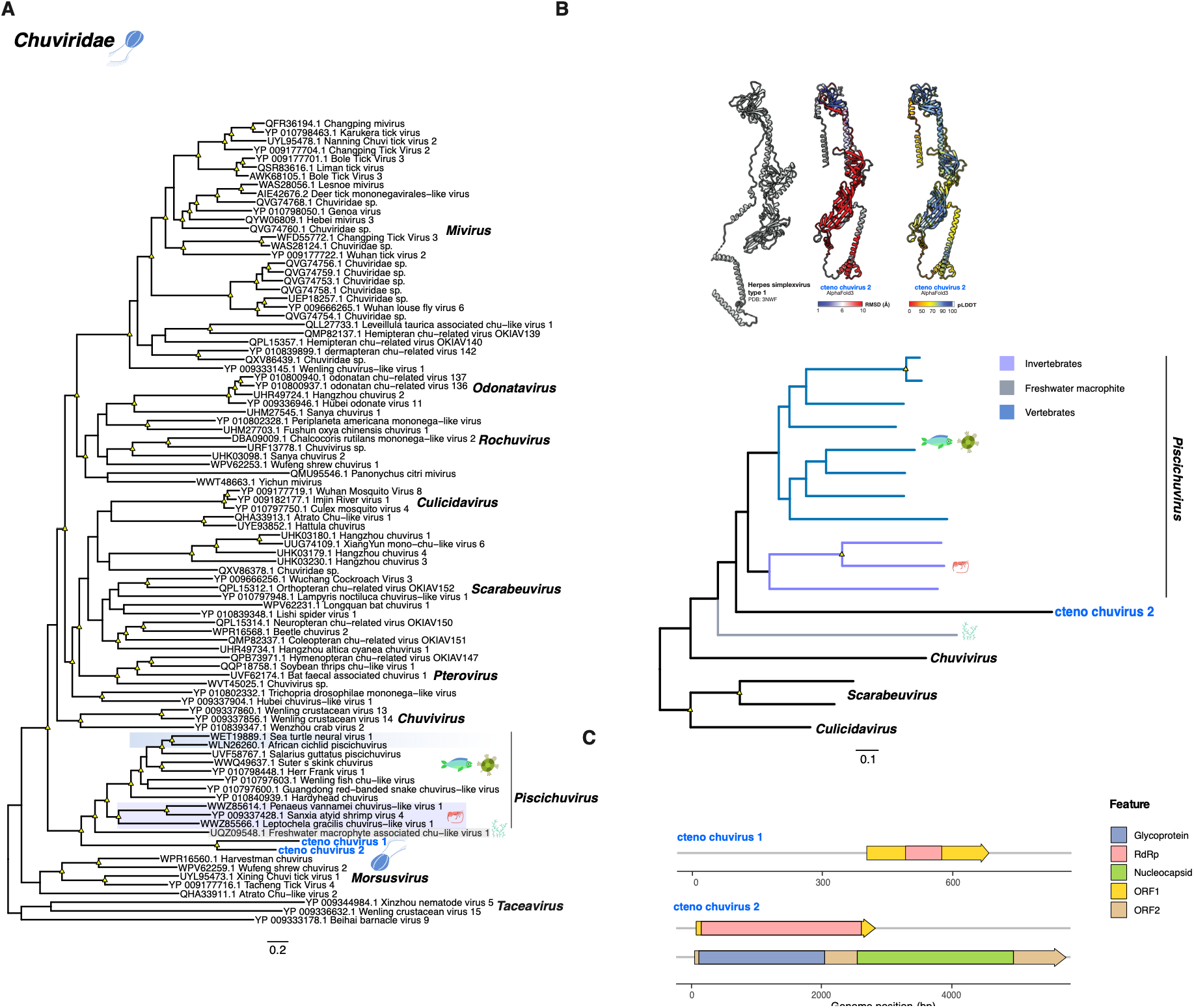
Phylogenetic relationships and conserved domains for the novel chuviruses identified in Ctenophora. (A) Phylogenetic placement of the novel cteno chuvirus as sister-group to the genus *Pisci*c*huvirus* based on the amino acid sequences of the RdRp. The viruses identified in this study are indicated in bold blue text. Newly discovered viruses are indicated by jelly comb silhouettes. The scale bar indicates the number of substitutions per site. (B) *Top panel*: Reference crystal structure of the Herpes simplex virus type 1 glycoprotein (class III fusion protein, PDB 3NWF), representing the closest homolog of the cteno chuvirus glycoprotein identified in Foldseek structural similarity searches of the RCSB Protein Data Bank (RCSB PDB). The predicted structure is coloured based on the RMSD and pLDDT scores as indicated in the bars. *Bottom panel*: Phylogenetic tree of the glycoprotein based on a partitioned model combining 3Di and amino acid alignment. The scale bar indicates the number of 3Di and amino acid character substitutions per site. The trees are midpoint rooted for purpose of clarity. Well-supported nodes are denoted with yellow triangle shapes (SH-aLRT ≥ 80 % and UFboot ≥ 95 %). (C) Schematic depiction of the ORFs and domains recovered for the novel cteno chuviruses.

## Discussion

Basal metazoan lineages potentially provide an important perspective on virus evolution. The unique biology of the ctenophores and placozoans has prompted research into their evolution, comparative genomics and developmental biology, in turn providing insights into their origins, physiology, and the development of different organs systems in animals [4, 24]. Despite this, the RNA viromes of ctenophores and placozoans remain poorly characterized, leaving a substantial gap in our understanding of the evolutionary history of RNA viruses [32–34]. Hence, investigating the diversity and phylogenetic relationships of RNA viruses in ctenophores and placozoans can shed light on their origin as well as broader patterns of virus-hosts associations in animals.

Our analysis of publicly accessible SRA data sets revealed a broad diversity of RNA viruses in ctenophores and placozoans, greatly expanding the host range of several virus groups. While our results revealed previously undetected RNA virus groups (e.g., chuviruses and endornaviruses), we also failed to recover others (e.g., mitoviruses and partitiviruses) previously identified in virome composition analyses of ctenophores and placozoans [32]. This likely reflects differences in methodological approaches from earlier work based on classification of sequencing reads [32]. In contrast, the current study relies on contig-based virus taxonomic classification and targeting of contigs containing RdRp sequences. [32]

Of particular note, with their detection in two ctenophore species we expanded the host range of the *Jingchuvirales* to early metazoans, and identified broader host shifts across multiple kingdoms spanning plants, fungi and animals. For example, endornaviruses (*Martellivirales*), which are typically found in plants, fungi and oomycetes, were also detected in placozoan libraries, thereby likely expanding their host range into animals (i.e., early-diverging metazoans). However, a caveat remains regarding confident host assignment due to the high proportion of unclassified sequences (up to 82%), limiting taxonomic resolution in these libraries. Likewise, the detection of narnaviruses (*Wolframvirales*) and rhabdoviruses (*Mononegavirales*) in ctenophores and placozoans suggest a long term association within Metazoa consistent with host switching events [65]. Similarly, the basal placement of the novel cteno lispi-like virus within the *Lispiviridae* indicates a long-standing association of these viruses with Metazoa, perhaps covering virus-host co-divergence over the entire evolutionary history of this group.

Although cteno orthomyxo-like virus was originally detected in a whole individual of *Mnemiopsis leidyi* (Ctenophora: Lobata), the detection of this virus in embryos is consistent with an early developmental establishment [66, 67]. In addition, that we identified mononegaviruses (*Lispiviridae* and *Rhabdoviridae*) and picornaviruses in additional embryonic libraries further suggest that viral infection is already underway at very early developing stages in ctenophores. This contrasts with early work on cnidarians, in which virus detection was confined to a single retrovirus in embryos [66]. Additional research is needed to better understand aspects such as the persistence of the viral infection across life stages, tissue tropism and their potential effects on ctenophore development and survival.

Of particular note was the detection of the novel cteno orthoflavi-like virus (∼11 Kb) in *Beroe ovata* (Ctenophora: Beroida), which grouped with the Beroe forskalii cigar comb jelly flavi−like virus (346 nt bp) to form a Ctenophora clade within the *Flaviviridae*. That both viruses were recovered from *Beroida* suggests that these flaviviruses might either have co-diverged with their hosts or undergone host-jumping facilitated by ecological overlap of these species [67]. Moreover, the presence of cteno orthoflavi-like virus at high abundance in the aboral organ, a densely neuralized region [4, 21], raises the question of whether the virus exhibits tissue specificity or occurs systemically. Interestingly, while the phylogenetic analysis of the RdRp places the orthoflavi-like virus with fish viruses, a different phylogenetic picture emerged from the partition model (i.e., the envelope glycoprotein sequence and structural data), where the novel Ctenophora lineage emerges as a sister branch to some vertebrate and invertebrate viruses, including fish viruses, indicating an early divergence relative to other lineages within the genus *Orthoflavivirus*. Because the exact placement of this lineage is sensitive to changes with the addition of 3Di (i.e., structural characters), the discovery of additional orthoflavi-like viruses and the continued maturation of structural phylogenetics methods will improve phylogenetic inference by breaking up long branches and enabling more accurate interpretations.

In a similar fashion, the discovery of the novel chuviruses in specimens from different ctenophore orders (i.e., Lobata and Cydippida) indicates the presence of a yet-unrecognized diversity of these viruses and a broader host range within ctenophores. Importantly, there is growing evidence that chuviruses are widespread among animals, occurring predominantly in invertebrates but also in some vertebrate hosts [68–72]. Based on the data generated here it seems plausible that piscichuviruses might have an origin tracing back to the early-diverging metazoans, followed by diversification into phylogenetically distant lineages that may have been facilitated by predator-prey interactions [73–75]. Sampling a broader range of taxa will be key to assessing this scenario [76]. Likewise, additional studies will be necessary to determine how trophic dynamics, receptor usage, and tissue tropism influence chuvirus transmission in ctenophores.

Although early diverging lineages metazoa might be expected to predominately harbour ancient viral lineages, only a subset of the ctenophore and placozoan viruses identified here occupied basal positions within their respective groups, particularly in ctenophores, reflecting a combination of long-term associations and host-jumping events. While some viruses occupied early divergent positions (i.e., *Chuviridae*, *Flaviriridae*, *Lispiviridae*, *Marnaviridae*), consistent with an early aquatic origin and subsequent diversification across multiple eukaryotic hosts, others likely reflect more recent cross-species transmission events mediated by host ecological interactions (i.e., *Endornaviridae*, *Mymonaviridae*, *Narnaviridae*, *Orthomyxoviridae*, *Picornavirales* (*Dicistroviridae* and unclassified), *Birnaviridae*, *Phasmaviridae*, *Orthototiviridae* and *Rhabdoviridae*). Indeed, for both the ctenophores and placozoans, the viruses observed tended to comprise distinct new lineages within established virus groups, with few occupying basal position. Notably, that some of the viruses identified here were connected to other taxa by long branches and had low support values prevents confident taxonomic assignments or conclusions regarding their evolutionary relationships to known virus sequences. For instance, despite the small sample size, rhabdoviruses and endornaviruses were detected at high abundance and formed distinct phylogenetic lineages, reflecting unsampled viral diversity and host-associated diversification in early metazoans [65]. Notably, the detection of a sequence identical to *Zymoseptoria tritici* fusarivirus 1 (100% id; *Durnavirales*) in placozoans suggests recent contamination from fungal sources. As an important caveat, although the use of scRNA-seq libraries increase confidence in host-virus association inference for placozoans, we cannot exclude the possibility that index-hopping or ambient viral sequences contributed to some assignments [77–79]. In addition, scRNA-seq has inherent limitations for capturing non-polyadenylated transcripts, potentially biasing detection for viruses lacking poly(A) tail such as flaviviruses and orthomyxoviruses [80].

Finally, this study identifies several important directions for future research. More work determining the extent of co-divergence and cross-species transmission in early-diverging metazoans will be critical to resolving host-virus associations and evolutionary histories. In addition, systematic searches for highly divergent viruses using homology-independent approaches will be essential to uncovering diversity that is undetected by sequence-similarity methods.

## Data availability

Details of the raw sequence reads obtained from public SRA data sets, including their SRA accession numbers, are provided in Table 1. Virus sequences identified in this study are available under NCBI/GenBank accessions XXXX–YYYYY.

## Acknowledgements

Data analysis was conducted using the computational resources available on the National Computational Infrastructure (NCI) and data resources provided by the Australian BioCommons Leadership Share (ABLeS) program. This program is co-funded by Bioplatforms Australia (enabled by NCRIS), the National Computational Infrastructure and Pawsey Supercomputing Research Centre.

## Supplementary Material

**Supplementary Figure S1.** ML phylogenetic trees based on the RdRp of virus groups found in ctenophores and placozoan libraries. The viruses identified in this study are denoted by coloured tip points, with blue indicating ctenophore-associated viruses and orange indicating placozoan-associated viruses. The scale bar indicates the number of substitutions per site. Well-supported nodes are denoted with yellow triangle shapes (SH-aLRT ≥ 80 % and UFboot ≥ 95 %). The trees are midpoint rooted for clarity.

**Supplementary Figure S2.** Glycoprotein structure modelling for the novel cteno orthoflavi-like virus. (A) Reference crystal structure of the Zika virus glycoprotein (PDB 7A3N) and the AlphaFold3-predicted model for cteno orthoflavi-like virus glycoprotein. The predicted structure is coloured based on the pLDDT and RMSD (based on superposition with PDB 7A3N) scores as indicated in the bars. The reference structure was selected based on structural homology searches of the PDB using Foldseek. (B) Foldseek structure-based similarity heat map comparing the predicted structures for cteno orthoflavi-like virus with experimentally determined glycoprotein structures from fusion protein classes I-III, with colour indicating E-value scores.

**Supplementary Figure S3.** ML phylogenetic trees based on the glycoprotein of viruses within the genus *Orthoflavivirus.* Trees were inferred using three approaches: (A) amino acid sequence alignment, (B) 3Di alignment and (C) Partitioned model combining both 3Di and amino acid sequence data. The scale bar indicates the number of substitutions per site. Strong node support (SH-aLRT ≥ 80% and UFboot ≥ 95%) is indicated by a black circle while intermediate support (SH-aLRT < 80% and UFboot > 95% or SH-aLRT > 80% and UFboot < 95%) is indicated by a half-filled circle. Clades are coloured as indicated in the key. The tree is rooted using nairoviruses as the outgroup. All unclassified orthoflaviviruses are in black, with the novel cteno orthoflavivirus in bold.

**Supplementary Figure S4.** Glycoprotein structure modelling for the novel cteno chuvirus. (A) Reference crystal structure of the Herpes simplexvirus virus glycoprotein (a class III fusion protein) and the AlphaFold3-predicted model for cteno chuvirus glycoprotein. The predicted structure is coloured based on the pLDDT and RMSD based on superposition with PDB 7A3N) scores as indicated in the bars. The reference structure was selected based on structural homology searches of the PDB using Foldseek. (B) Foldseek structure-based similarity heat map comparing the predicted structures for cteno chuvirus with experimentally determined glycoprotein structures from fusion protein classes I-III, with colour indicating E-value scores.

**Supplementary Figure S5.** ML phylogenetic trees based on the glycoprotein of viruses within the family *Chuviridae.* Trees were inferred using three approaches: (A) amino acid sequence alignment, (B) 3Di alignment and (C) Partitioned model combining both 3Di and amino acid sequence data. Strong node support (SH-aLRT ≥ 80% and UFboot ≥ 95%) is indicated by a black circle while intermediate support (SH-aLRT < 80% and UFboot > 95% or SH-aLRT > 80% and UFboot < 95%) is indicated by a half-filled circle. Clades are coloured as indicated in the key. All piscichuvirus-like sequences are in light blue, with the novel cteno chuvirus 2 sequence in bold.

**Supplementary Table S1.** AlphaFold and Foldseek structural analyses for the predicted orthoflavi-like and chuvi-like viruses. This excel file contains three worksheets. (i) Foldseek_references: Experimental PDB structures from viral membrane-fusion protein classes I–III used for querying and validation of the predicted glycoprotein structures for the novel flavivirus and chuvirus sequences. (ii) Foldseek_results: contains the Foldseek structural similarity searches results between predicted viral glycoproteins and the reference fusion proteins. (iii) Structural_prediction_metrics: summarises AlphaFold prediction confidence metrics for each predicted glycoprotein structure.

**Supplementary Table S2.** Amino acid and 3Di substitution models chosen by ModelFinder for the phylogenetic analyses of the flavivirus and chuvirus glycoproteins.

