## Supplementary material for "The RNA virome of early metazoans sheds light on long-term virus-host relationships": Figure S1

#### *Picornavirales*

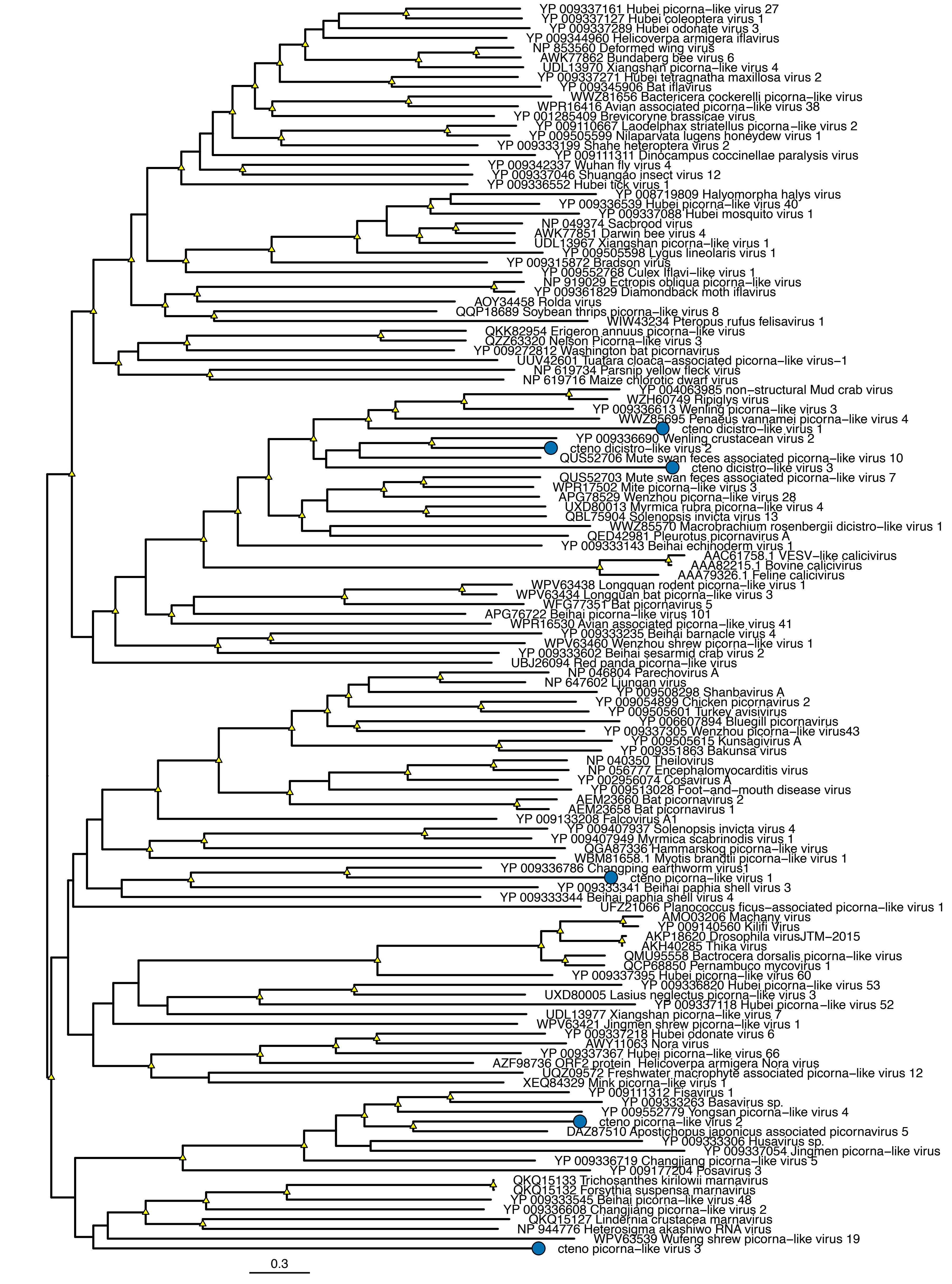

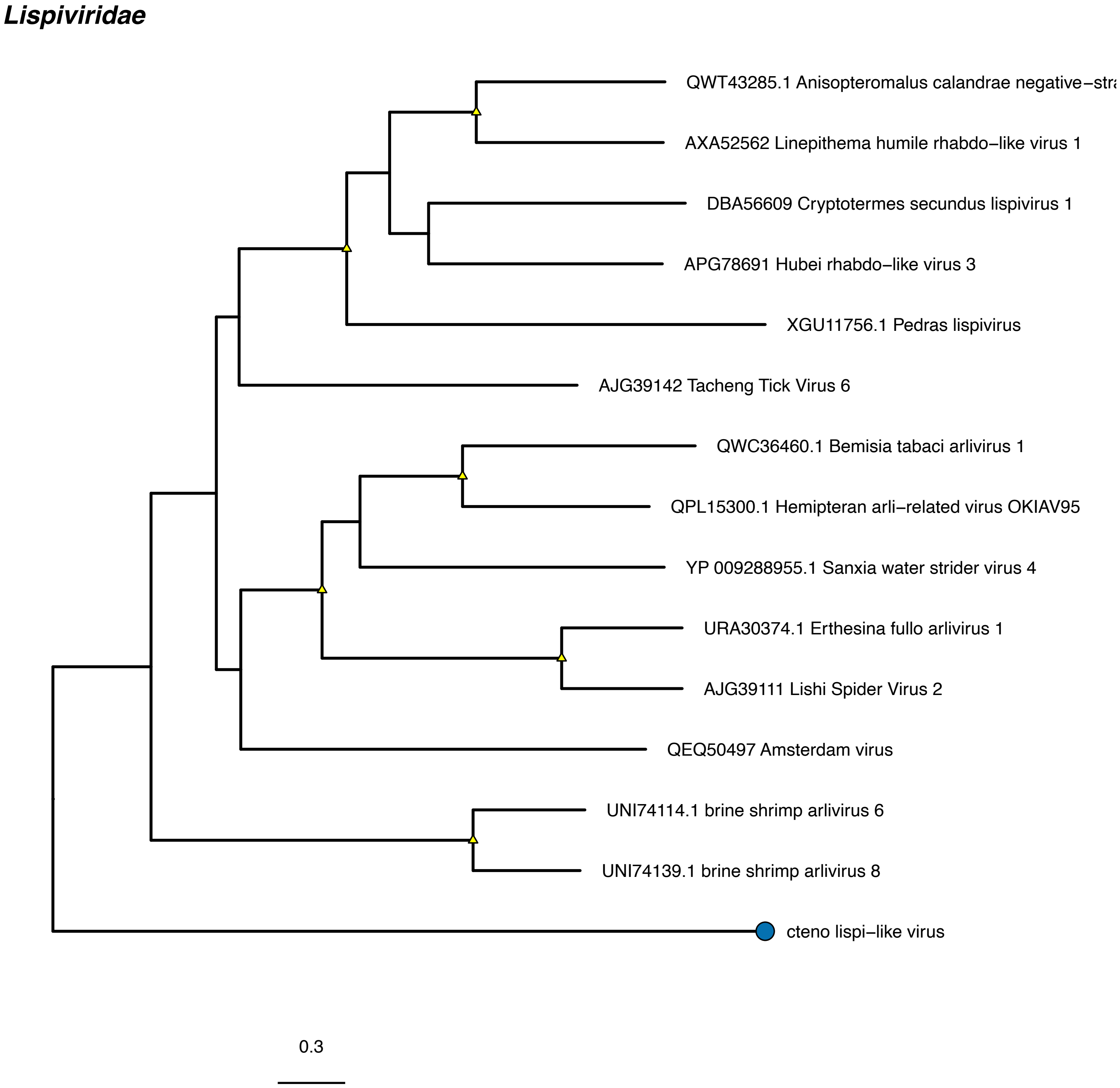
Lispiviridae
QWT43285.1 Anisopteromalus calandrae negative−stra
AXA52562 Linepithema humile rhabdo−like virus 1
DBA56609 Cryptotermes secundus lispivirus 1
APG78691 Hubei rhabdo−like virus 3
XGU11756.1 Pedras lispivirus
AJG39142 Tacheng Tick Virus 6
QWC36460.1 Bemisia tabaci arlivirus 1
QPL15300.1 Hemipteran arli−related virus OKIAV95
YP 009288955.1 Sanxia water strider virus 4
URA30374.1 Erthesina fullo arlivirus 1
AJG39111 Lishi Spider Virus 2
QEQ50497 Amsterdam virus
UNI74114.1 brine shrimp arlivirus 6
UNI74139.1 brine shrimp arlivirus 8
cteno lispi−like virus
0.3

***Orthomyxoviridae***

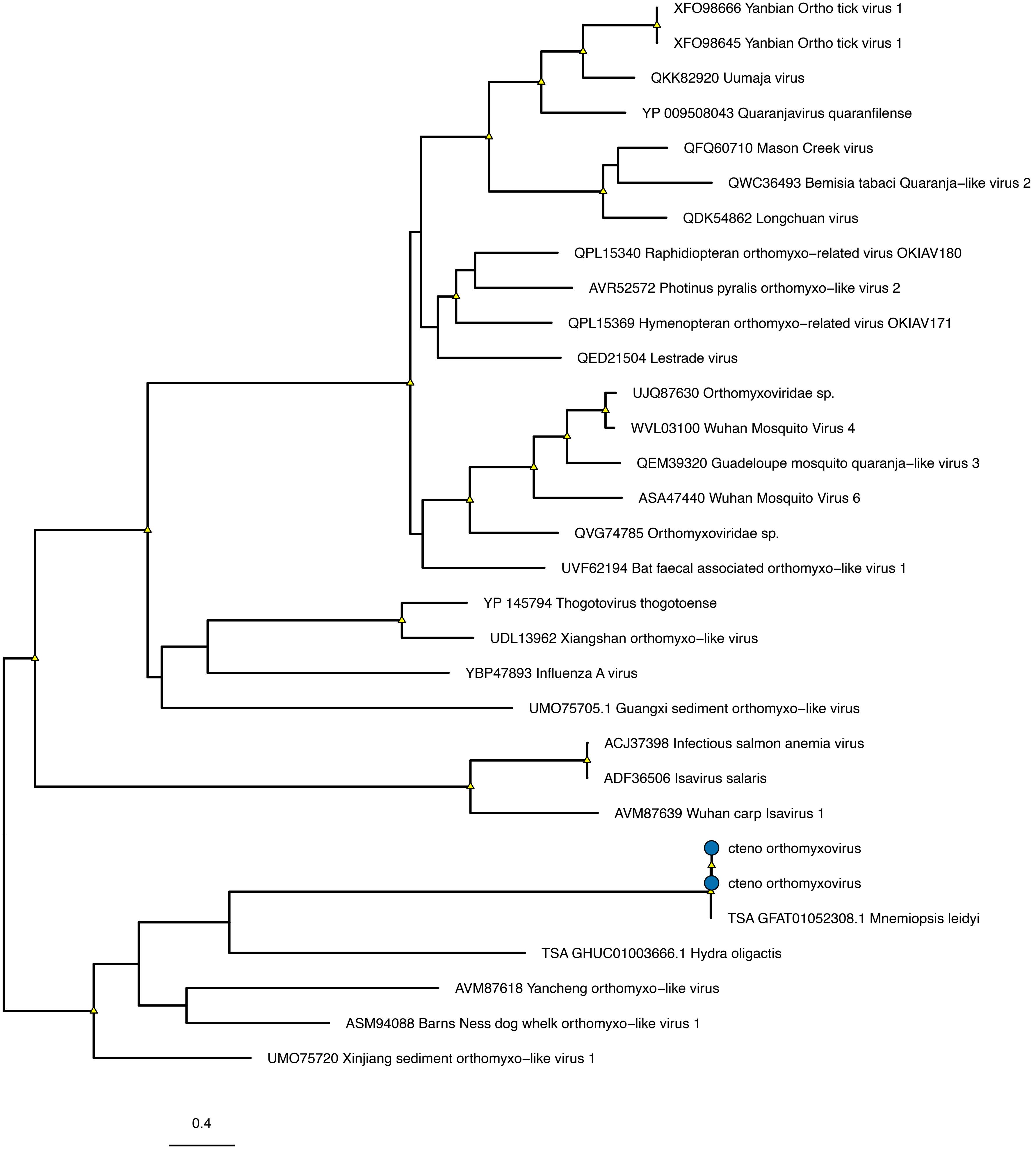

### *Orthototiviridae*

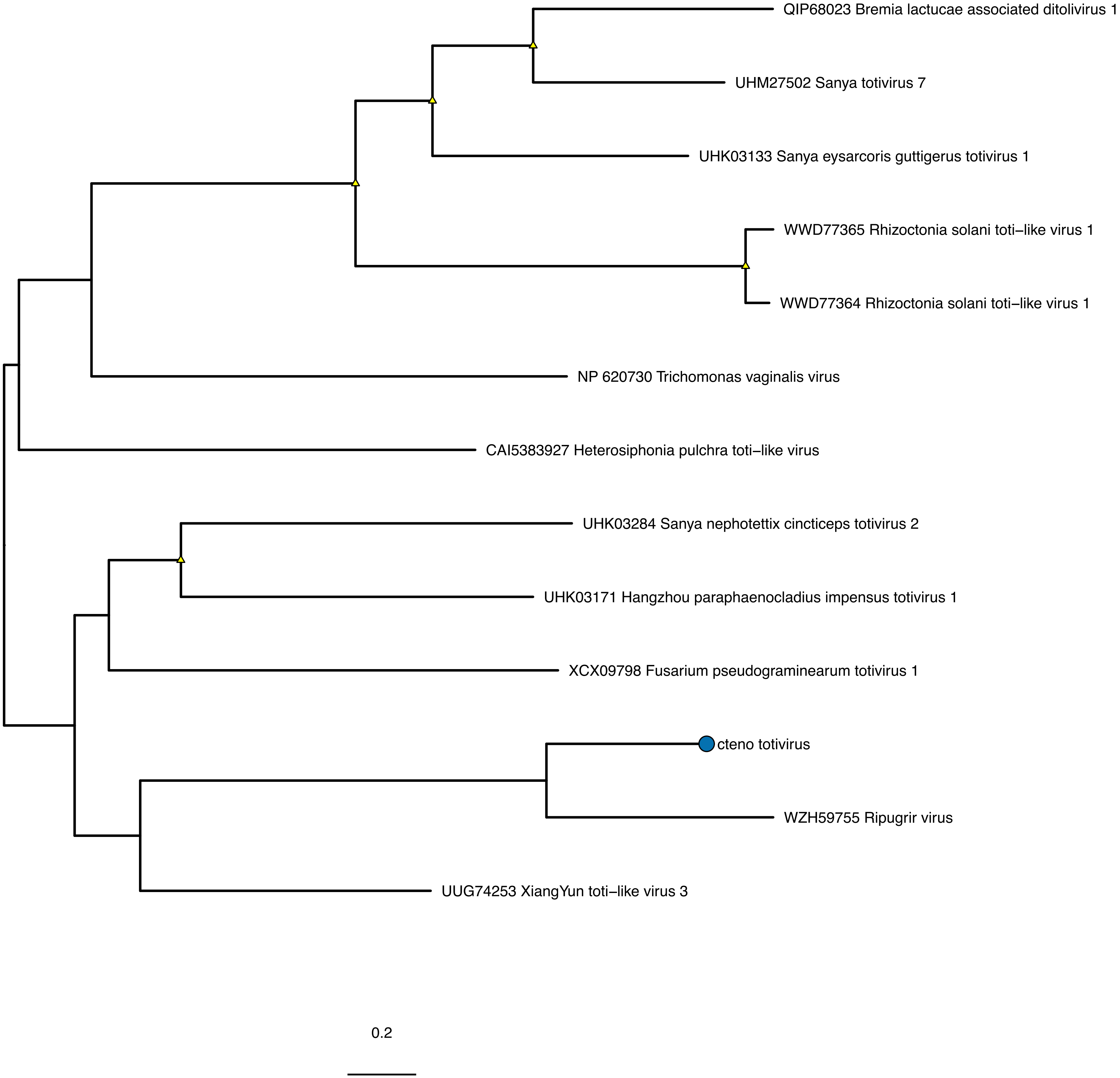

#### *Birnaviridae*

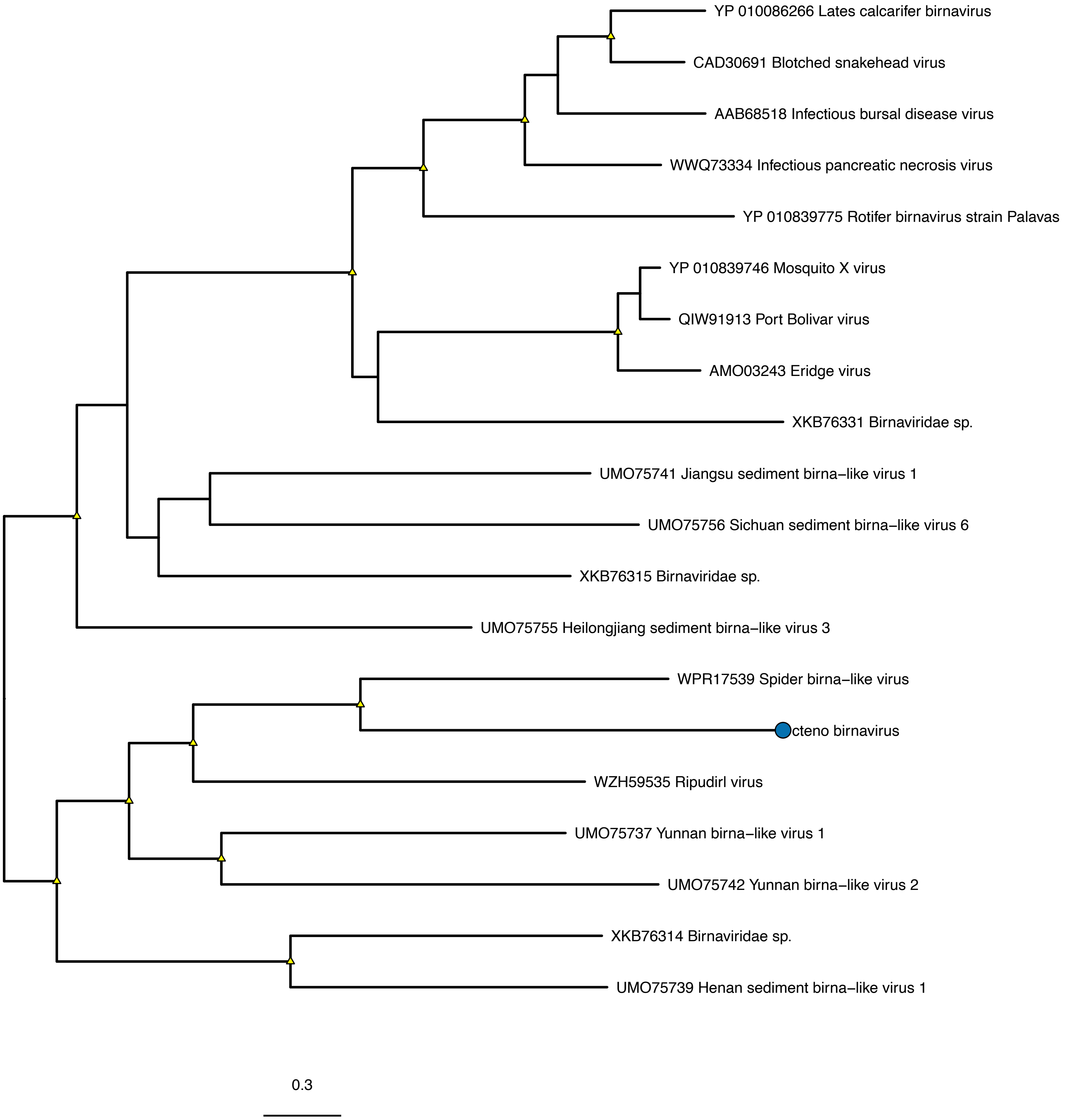

#### *Phasmaviridae*

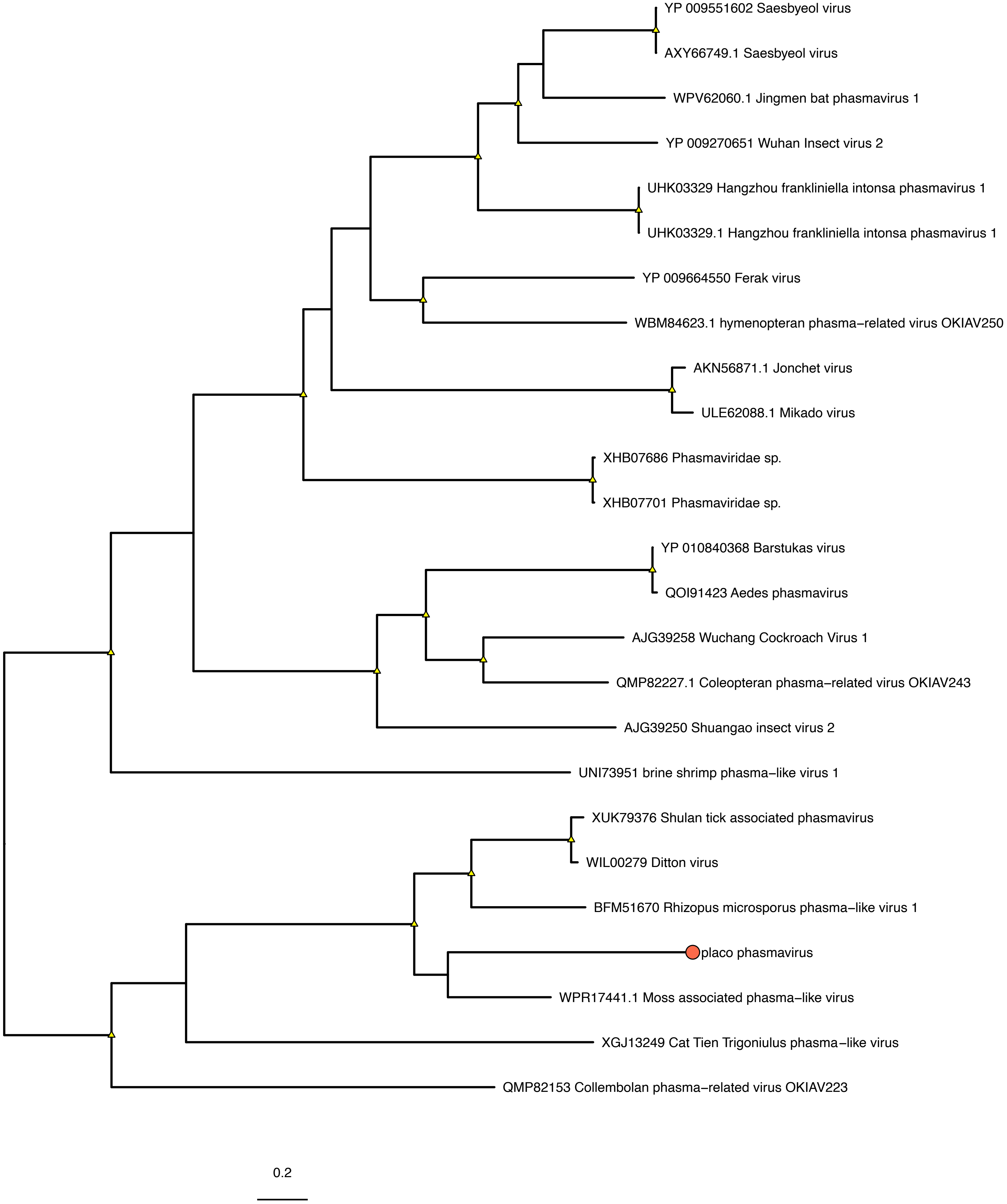

#### *Fusariviridae*

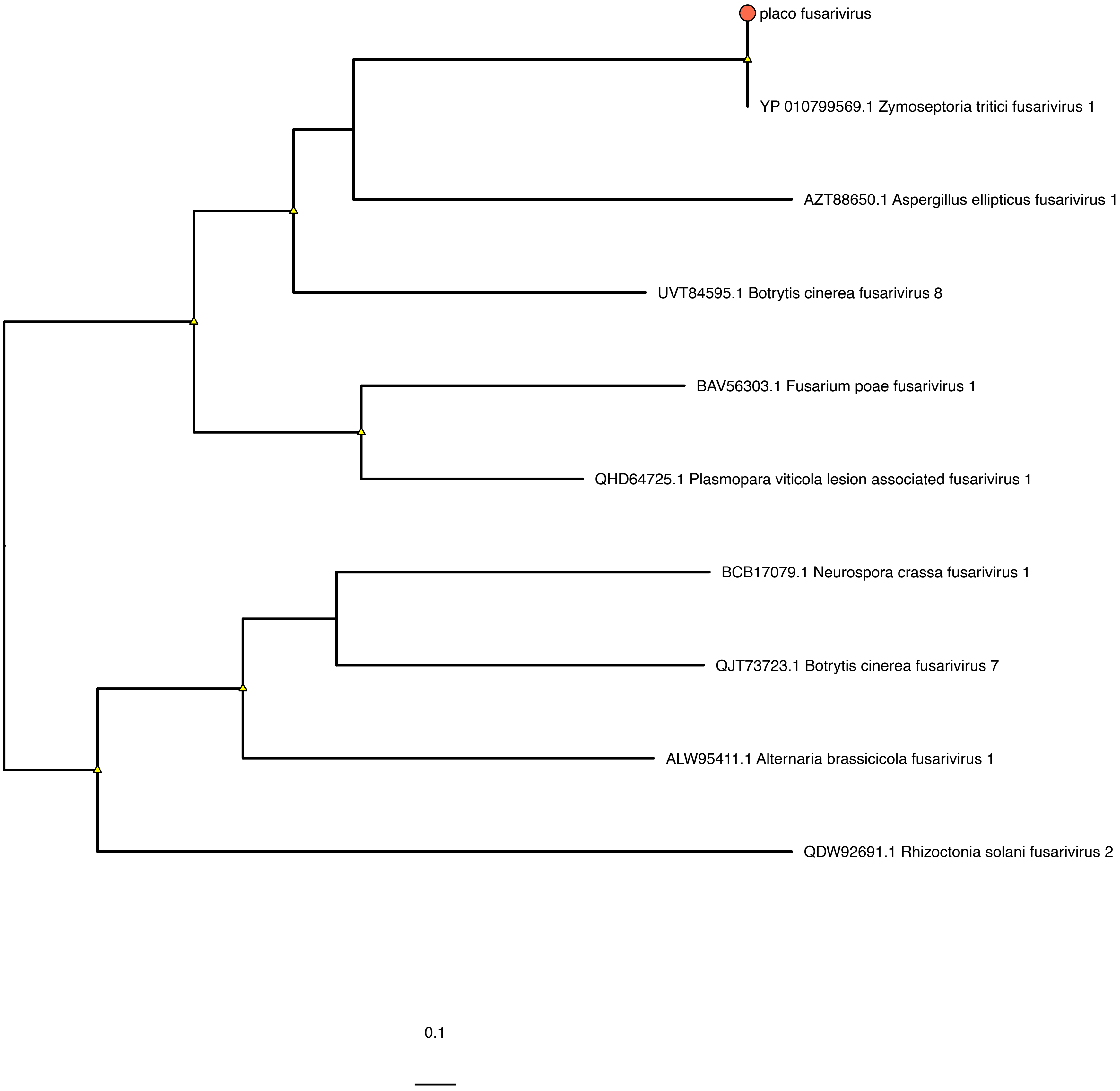

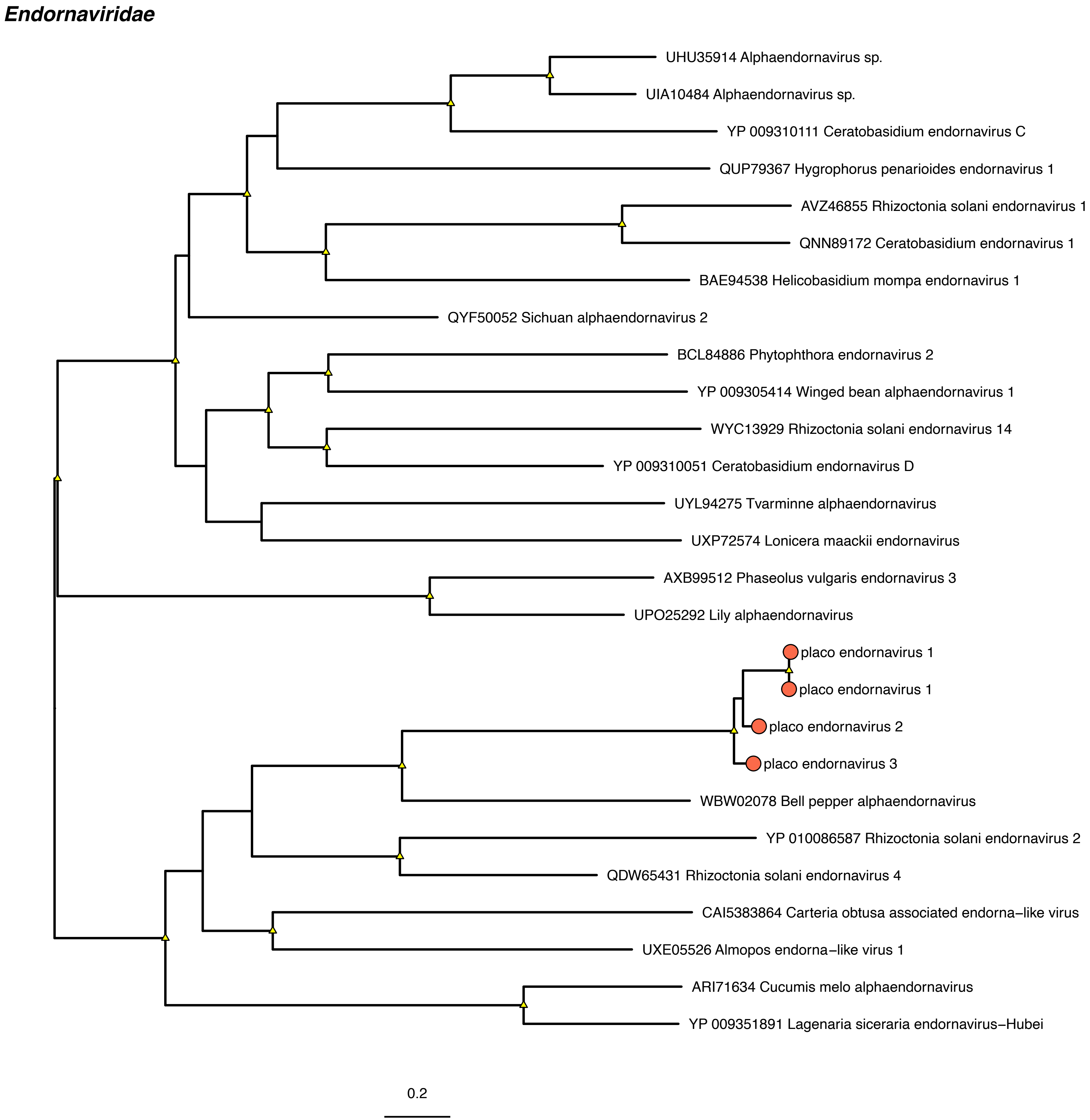
Endornaviridae
UHU35914 Alphaendornavirus sp.
UIA10484 Alphaendornavirus sp.
YP 009310111 Ceratobasidium endornavirus C
QUP79367 Hygrophorus penarioides endornavirus 1
AVZ46855 Rhizoctonia solani endornavirus 1
QNN89172 Ceratobasidium endornavirus 1
BAE94538 Helicobasidium mompa endornavirus 1
QYF50052 Sichuan alphaendornavirus 2
BCL84886 Phytophthora endornavirus 2
YP 009305414 Winged bean alphaendornavirus 1
WYC13929 Rhizoctonia solani endornavirus 14
YP 009310051 Ceratobasidium endornavirus D
UYL94275 Tvarminne alphaendornavirus
UXP72574 Lonicera maackii endornavirus
AXB99512 Phaseolus vulgaris endornavirus 3
UPO25292 Lily alphaendornavirus
placo endornavirus 1
placo endornavirus 1
placo endornavirus 2
placo endornavirus 3
WBW02078 Bell pepper alphaendornavirus
YP 010086587 Rhizoctonia solani endornavirus 2
QDW65431 Rhizoctonia solani endornavirus 4
CAI5383864 Carteria obtusa associated endorna−like virus
UXE05526 Almopos endorna−like virus 1
ARI71634 Cucumis melo alphaendornavirus
YP 009351891 Lagenaria siceraria endornavirus−Hubei
0.2

#### *Rhabdoviridae*

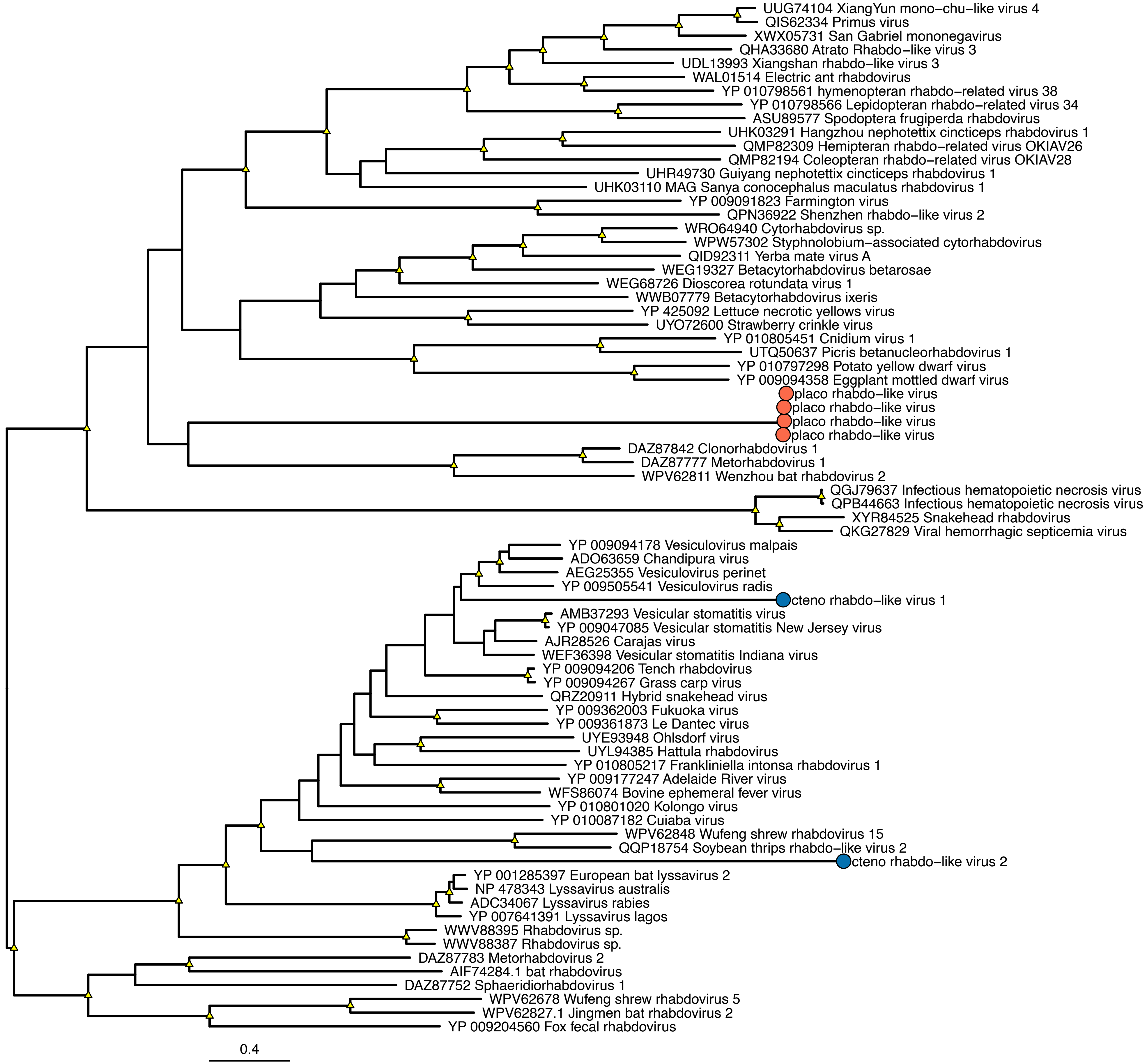

*Mymonaviridae*

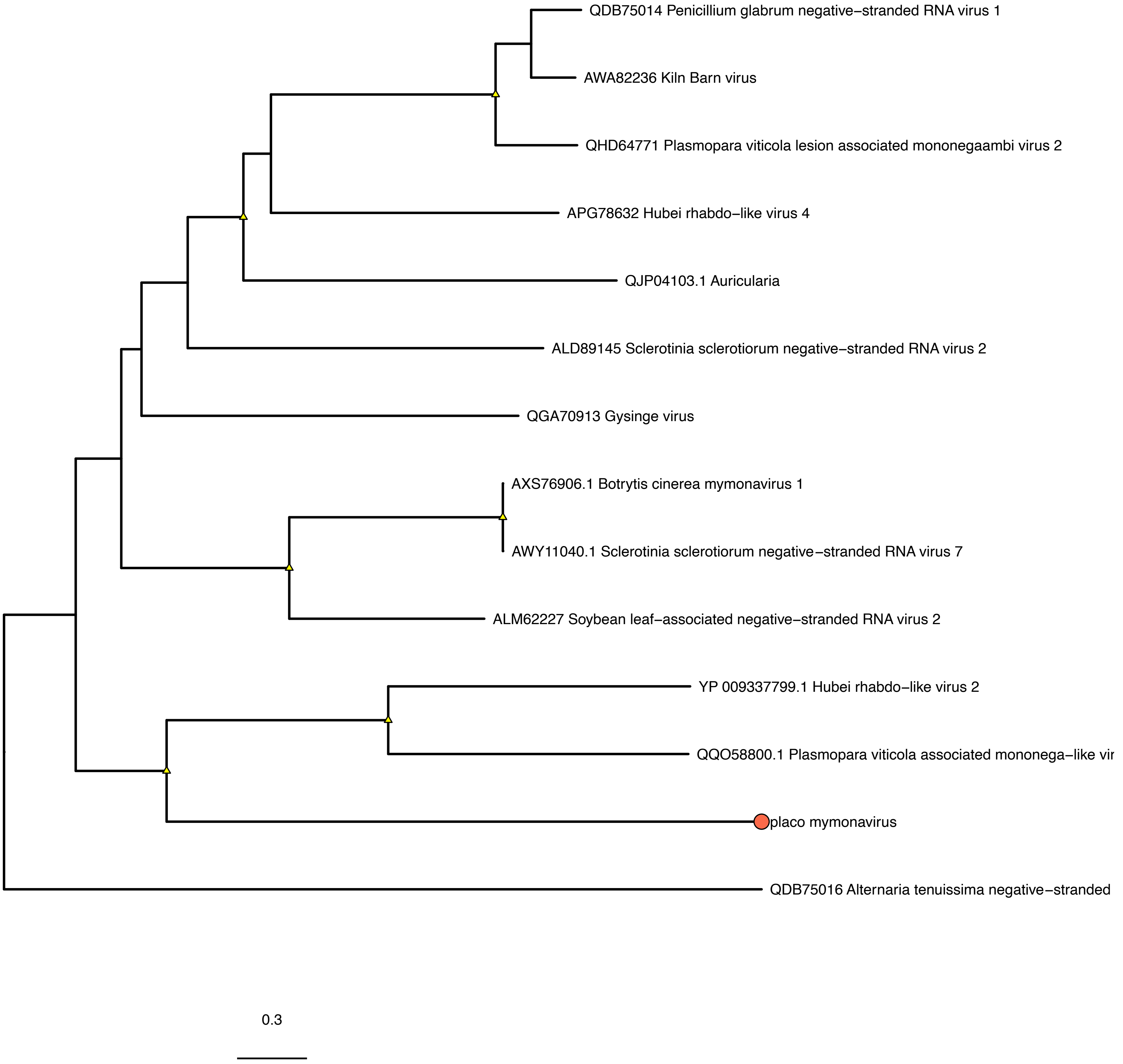

***Narnaviridae***

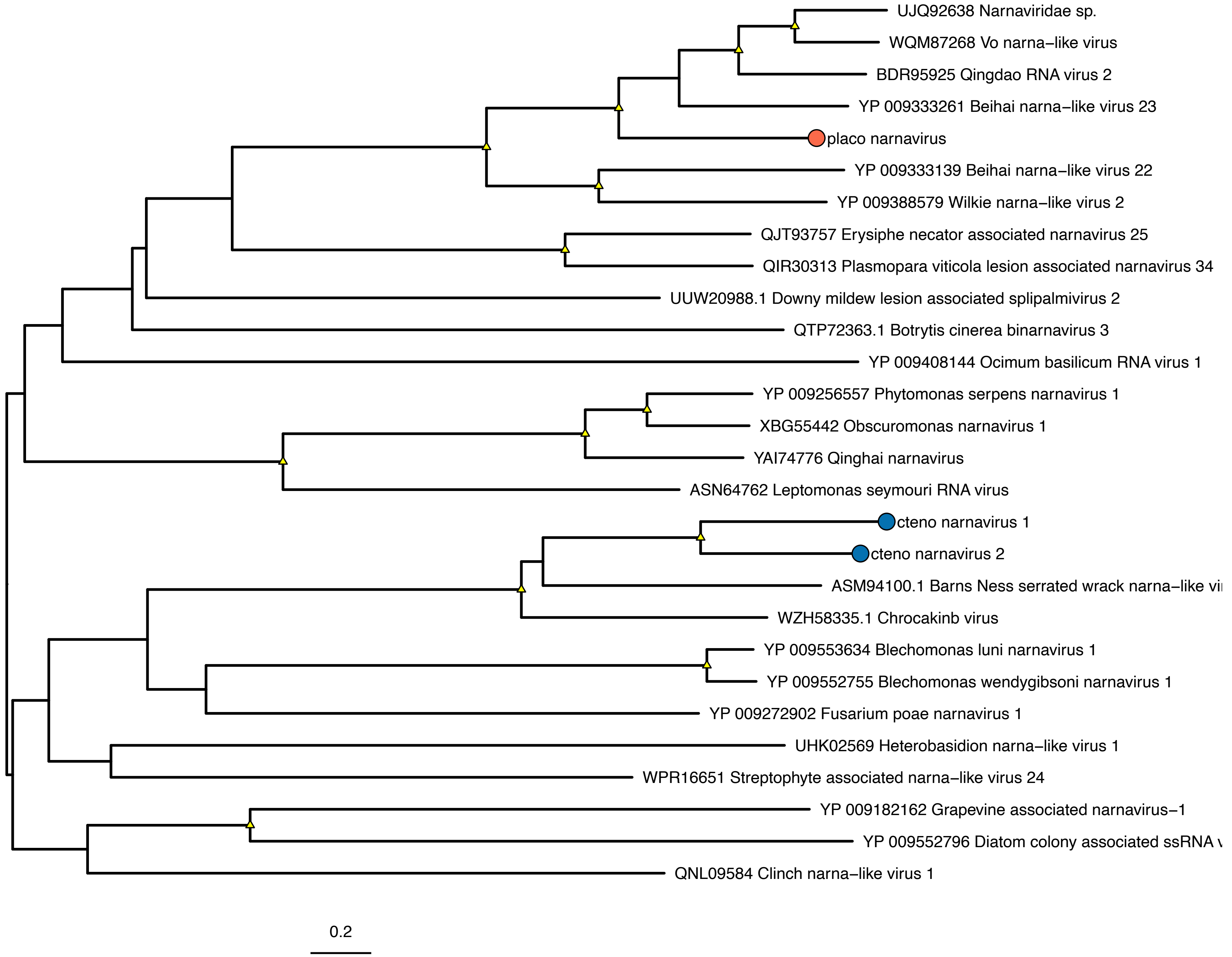
