## Supplementary figures and images for "The RNA virome of early metazoans sheds light on long-term virus-host relationships"

### Figure S2

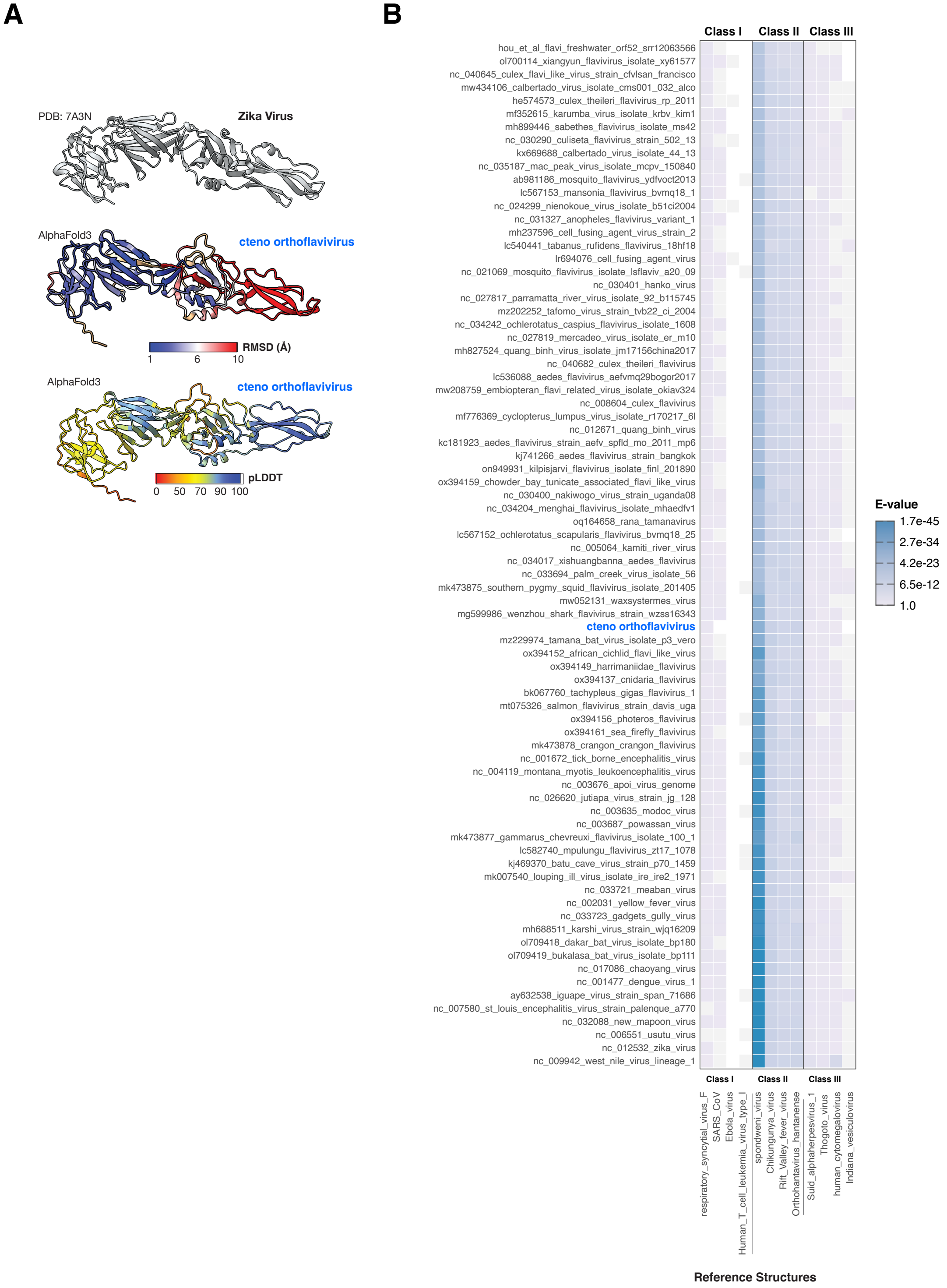

### Figure S3

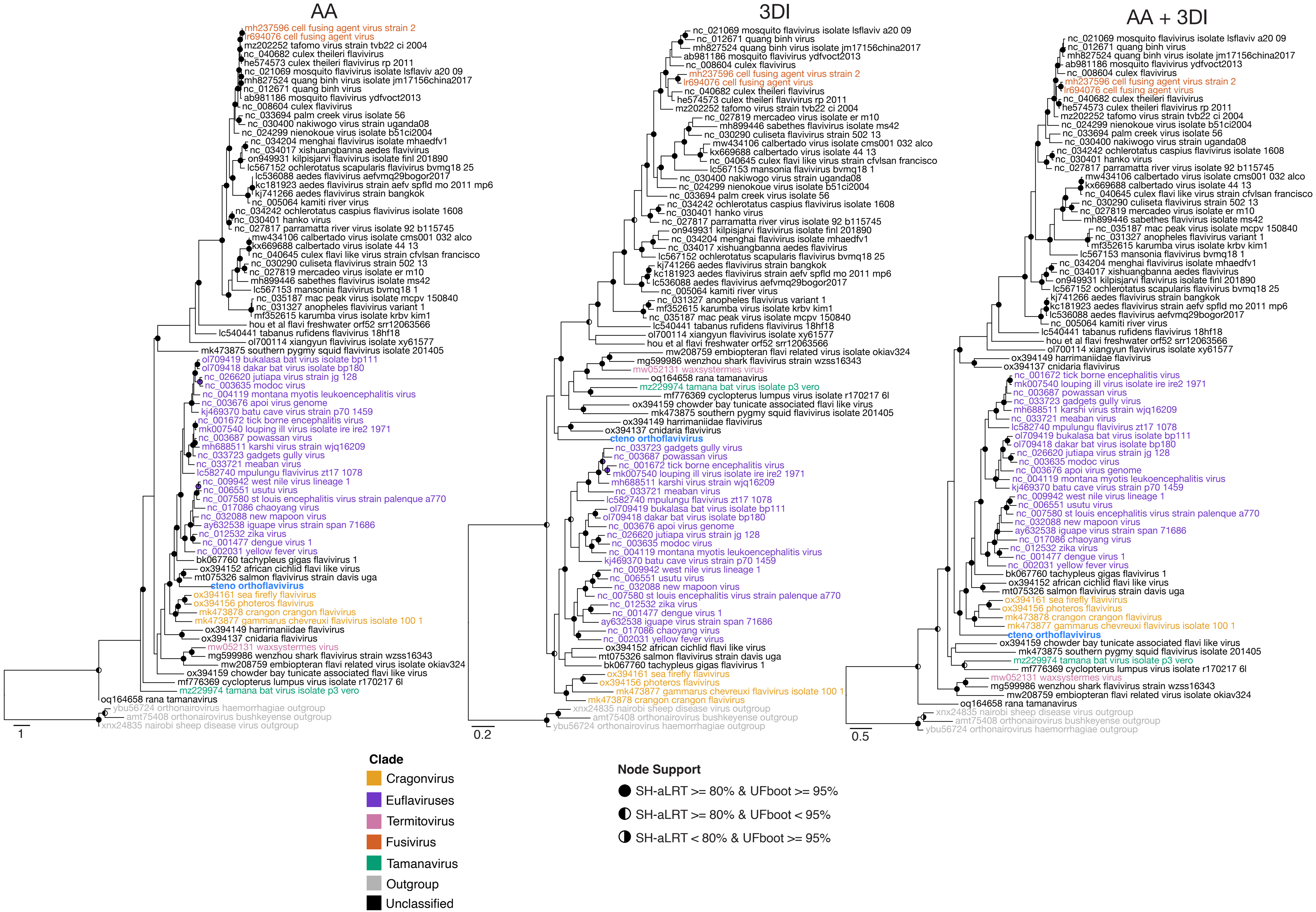

### Figure S4

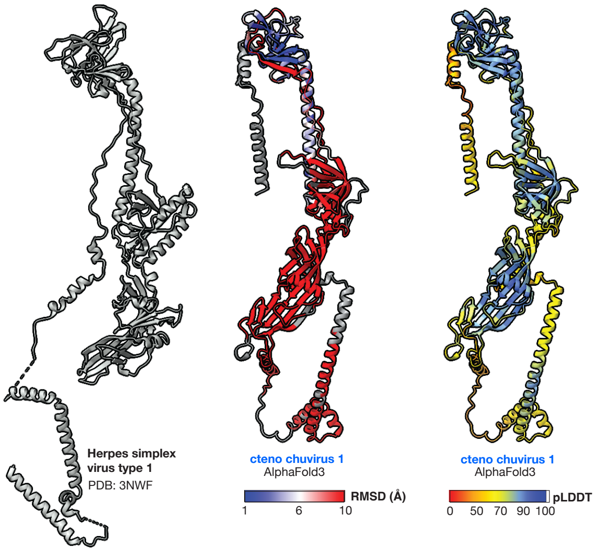

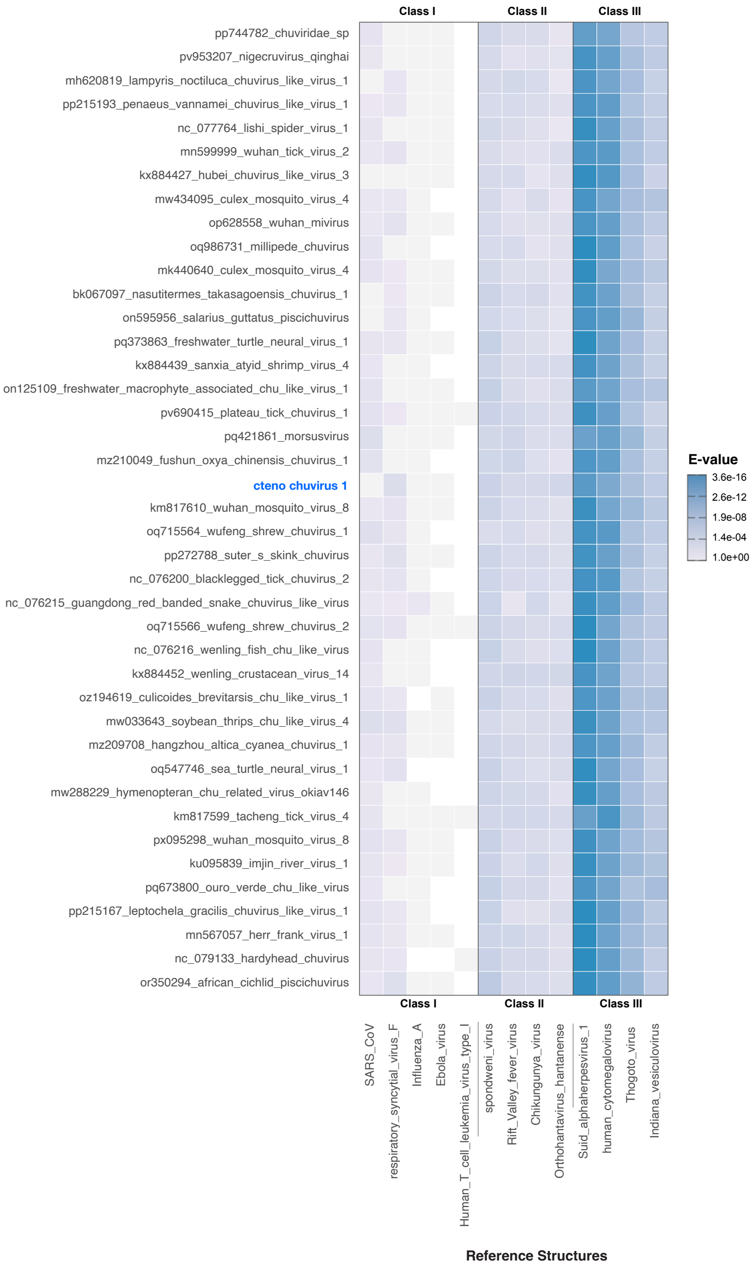

### Figure S5

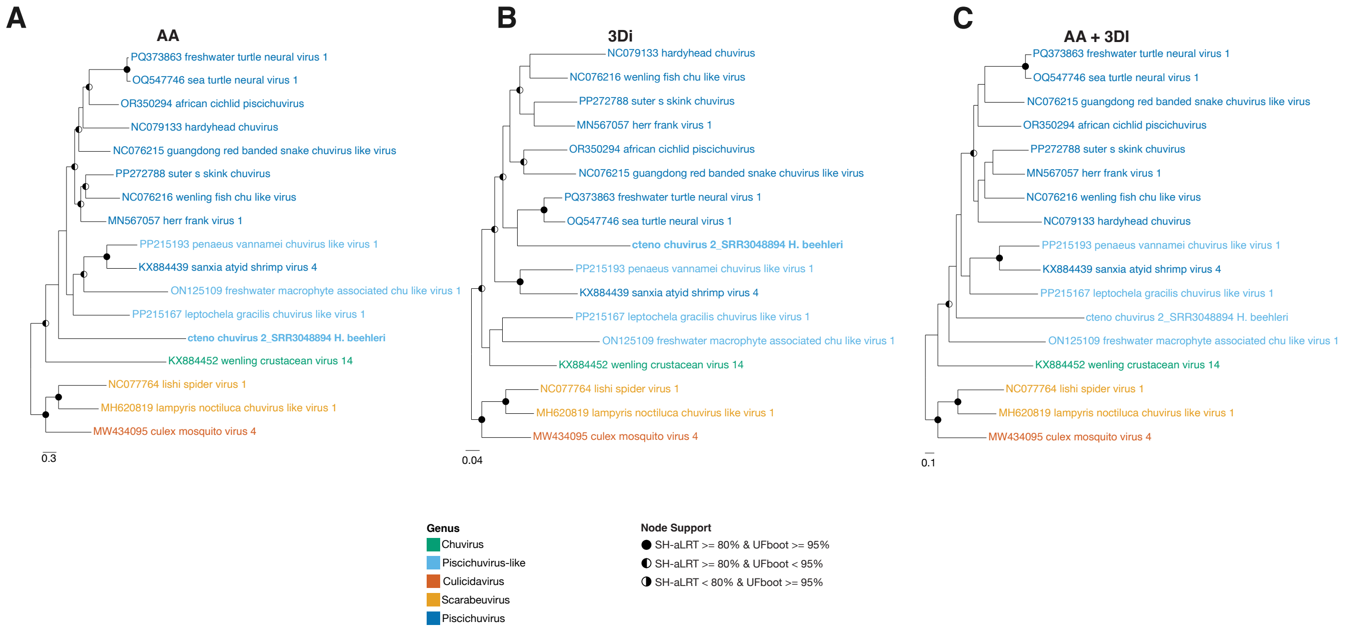
